# Gα_q/11_ signalling counteracts endothelial dysfunction in the brain and protects cognition in aged mice

**DOI:** 10.1101/2025.07.24.666499

**Authors:** Dimitrios Spyropoulos, Lena Kleindienst, Phillip Ehrich, Dorothea Ziemens, Rentsenkhand Natsagdorj, Nicholas Noll, Cathrin Elly Hansen, Géza Adam Curley, Anne-Sophie Gutt, Ákos Menyhárt, Jan Sedlacik, Peter Ludewig, Beate Lembrich, Ümit Özorhan, Evi Kostenis, Jens Fiehler, Gereon Hüttmann, Thomas Andrew Longden, Ruben Nogueiras, Vincent Prevot, Helge Müller-Fielitz, Stefan Offermanns, Nina Wettschureck, Eszter Farkas, Markus Schwaninger, Jan Wenzel

## Abstract

**Background:** Cerebral small vessel disease (cSVD) is a major cause of stroke and dementia, and is associated with increased blood-brain barrier permeability, neuroinflammation, and endothelial dysfunction. Endothelial Gα_q/11_ proteins are involved in vascular tone regulation and have been shown to affect capillary blood flow in the brain. Since factors downstream of activated Gα_q/11_ proteins, such as endothelial NO synthase (eNOS) activity are discussed in cSVD, we wondered whether the brain endothelial Gα_q/11_ signalling pathway might influence cSVD-related pathology.

**Methods:** Here, we generated mice carrying a brain endothelial-specific deletion of the Gα_q/11_ signalling and characterised these mice using different imaging and staining techniques, as well as behaviour tests measuring cognition in adult and aged mice. Immunoblots, electrophysiology, perfusion measurements, and *in vitro* experiments complemented those techniques.

**Findings:** The brain endothelial Gα_q/11_ signalling pathway preserves normal vascular reactivity, and its loss resembles mild endothelial dysfunction in the brain. While the vessel structure was maintained in adult mice, a deletion of the Gα_q/11_ signalling led to capillary rarefaction and blood-brain barrier disruption in aged mice. These effects were accompanied by disturbed VEGF signalling and an increase in senescence markers and oxidative stress in the vasculature, culminating in cognitive impairment with increased tau phosphorylation in the cortex and hippocampus, and decreased myelination in the white matter.

**Interpretation:** These findings reflect the main hallmarks of cSVD and demonstrate a protective role of Gα_q/11_ in endothelial cells in ageing. Furthermore, our results show that the combination of cerebral endothelial dysfunction and ageing accelerates cognitive impairment.

**Funding:** Research was supported by grants from the European Research Council (2019-WATCH-810331 to R.N., V.P., and M.S.), the Deutsche Forschungsgemeinschaft (SCHW 416/12-1 to M.S.; WE 6456/1-1 to J.W.; GRK1957 to M.S., H.M.-F., and J.W.), the institutional priority program MI-VascAD of the University of Lübeck (to H.M.-F. and J.W.), and the Marie-Sklodowska-Curie European Union’s Horizon 2020 research program (ENTRAIN-813294 to M.S. and J.W.).

**RESEARCH IN CONTEXT:** *Evidence before this study:* Cerebral small vessel disease (cSVD) is associated with advanced age, hypertension, and metabolic diseases. It is considered causal for a substantial part of stroke and dementia cases. Endothelial dysfunction is described as being present in all of the diseases mentioned, and it is discussed to play a role in the development of cSVD. However, most of the evidence is correlational rather than causal, and studies are usually carried out on subjects with an underlying disease for which endothelial dysfunction is a side effect.

*Added value of this study:* Here, we describe a novel brain endothelial signalling pathway that protects against endothelial dysfunction and loss of cerebrovascular reactivity. Without Gα_q/11_ signalling exclusively in the brain endothelium, mice show an impaired ability to increase brain perfusion in response to vasodilatory stimuli, such as neuronal activity. These mice, combined with an older age, develop vessel rarefaction and cognitive dysfunction, which can be explained by increased tau phosphorylation and white matter alteration.

*Implications of all the available evidence:* These findings, in the context of the existing studies, suggest that endothelial dysfunction *per se* is sufficient to induce cognitive impairment. They imply that the diagnoses and treatments for diseases associated with cognitive impairment should focus more on a dysfunctional endothelium. The Gα_q/11_ signalling pathway offers a new approach to address treatment endeavours in cSVD and, more broadly, in all diseases associated with a higher risk of vascular dementia and stroke.

## INTRODUCTION

Vascular impairment in the central nervous system is implicated in the development of cognitive dysfunction in the elderly, and several systemic diseases are associated with an increased risk of cognitive and vascular deficits.^1,2^ Cerebral small vessel disease (cSVD) is considered to be a leading cause of dementia and lacunar stroke.^3,4^ There are two main types of cSVD, the amyloid form with amyloid angiopathy, and the non-amyloid form, usually associated with old age, hypertension, diabetes and metabolic syndrome.^2^ In both cases, cSVD is characterised by white matter alterations, cerebral microinfarcts and bleeding, a disturbed blood-brain barrier (BBB), and cognitive impairment.^2^ The cerebral vasculature and especially capillary endothelial cells undergo profound changes during ageing,^5,6^ and endothelial dysfunction is strongly associated with cSVD. Still, the role of cerebral endothelial dysfunction in causing pathological changes in cSVD has not yet been sufficiently deciphered.^4^ Endothelial dysfunction refers to an impaired ability of the vasculature to respond to vasodilatory stimuli due to pathological alterations in the endothelium, which are strongly linked to reduced nitric oxide (NO) availability. NO production in the vasculature is mainly driven by the endothelial NO synthase (eNOS), which can be directly impaired or uncoupled in endothelial dysfunction.^7^ Interestingly, the partial eNOS-deficient mice are considered a model for cSVD and show hippocampal-dependent memory deficits, specifically at older ages.^8^ However, since eNOS germline deficiency leads to hypertension,^9^ the effect on the brain can be indirectly caused by peripherally-induced blood pressure cues.

eNOS activity is driven by phosphorylation and by an increase in intracellular calcium concentration.^7^ Both of these factors are strongly induced by the activation of Gα_q/11_ proteins. Gα_q/11_ proteins are activated by G protein-coupled receptors and lead to phospholipase C activation, which leads to depletion of phosphatidylinositol 4,5-bisphosphate (PIP_2_). This process not only leads to increased intracellular calcium via the generated inositol 3-phosphate (IP_3_) but also decreases PIP_2_ concentration. PIP_2_ can modulate electrical signalling in brain endothelial cells via interacting with ion channels ^10–12^ and was recently shown to restore perfusion in a cSVD mouse model.^13^ In addition, protein kinases like the protein kinase C (PKC), or AKT, are induced by Gα_q/11_ proteins and interfere directly with eNOS phosphorylation.

The deletion of the Gα_q/11_ pathway in all endothelial cells results in hypertension,^14^ whereas deletion only in the brain endothelium does not alter systemic blood pressure.^15^ However, how local endothelial function in the brain is affected by endothelial Gα_q/11_ is not clear. Altogether, these findings make the Gα_q/11_ signalling pathway in the brain endothelium an interesting target in the development of vascular-based cognitive dysfunction in cSVD.

Here, we demonstrate that the brain endothelial Gα_q/11_ signalling pathway is involved in maintaining normal vascular reactivity without profound structural and morphological changes in the braińs vasculature in adult mice. However, during ageing, mice lacking brain endothelial Gα_q/11_ (Gα_q/11_^beKO^) develop vascular rarefaction and increased blood-brain barrier integrity, leading to increased tau phosphorylation, decreased myelination in the white matter, and mild cognitive dysfunction, implying a causative role of brain endothelial dysfunction in ageing-associated cognitive deficits and cSVD.

## METHODS

### Mice

All the mouse lines were established on a C57BL/6 background. We used littermate mice that were sex- and age-matched between experimental groups. Mice of both sexes were used. The mice were kept at a constant temperature and humidity (22°C and 50%, respectively) on a 12-hour light/ dark cycle and were provided with standard laboratory chow and water *ad libitum*. All animal experiments were approved by the local animal ethics committee (Regierungspräsidium Karlsruhe; Ministerium für Landwirtschaft, ländliche Räume, Europa und Verbraucherschutz, Kiel, Germany) and were carried out according to the ARRIVE guidelines and to the EU Directive 2010/63/EU for animal experiments. Brain endothelial-specific knockout (beKO) animals were generated by using the BAC-transgenic *Slco1c1*-CreER^T2^ strain,^16^ which expresses the tamoxifen-inducible CreER^T2^ recombinase under the control of the mouse *Slco1c1* regulatory sequences in brain endothelial cells. The Cre mice were crossed with mice carrying a floxed Gα_q_ allele (*Gnaq*^fl/fl^), and they were Gα_11_-deficient (*Gna11*^-/-^)^17^ (*Gnaq*^fl/fl^; *Gna11*^-/-^), leading to a deletion of the Gα_q/11_ pathway in brain endothelial cells (Gα_q/11_^beKO^) after treatment with tamoxifen. Littermates lacking the Cre transgene and heterozygous for Gα_11_ (*Gnaq*^fl/fl^; *Gna11*^-/+^) were used as controls and received tamoxifen in parallel with the knockout animals. Deletion of both Gna11 and Gnaq is essential since these proteins can replace each other, and a single knockout is not sufficient to delete the whole pathway. In addition, in control mice, heterozygous *Gna11*-deleted mice are not expected to have any phenotype and were used as controls due to optimised breeding strategies, prioritising littermate experiments. To induce the deletion of the Gα_q/11_ pathway in astrocytes, we used a Glast-CreER^T2^ mouse line (Gα_q/11_^astroKO^),^18^ and to assess recombination in the *Slco1c1*-CreER^T2^ strain, we used the Ai14^+/fl^ reporter mouse, leading to tdTomato expression upon recombination.^19^ All animals were injected with 1 mg tamoxifen (Sigma-Aldrich) per 20 g body weight dissolved in 90% miglyol^®^ 812 /10% ethanol i.p. every 12 h for 5 consecutive days. AAV-mediated transduction of brain endothelial cells was achieved by using a specialised AAV capsid developed earlier.^20^ Gα_q/11_^BR1-beKO^ mice were generated by injecting 1×10^12^ genomic particles per mouse containing either a AAV-BR1 encoding a *CAG-Cre-2A-GFP* construct (i.v. injected into *Gnaq*^fl/fl^; *Gna11*^-/-^ mice) or a *CAG-GFP* construct (injected into *Gnaq*^fl/fl^; *Gna11*^-/+^ mice). *Cdh5*-GCaMP8 mice were described earlier,^21,22^ and were used to measure intracellular calcium activity in primary brain endothelial cells.

The investigators were blinded to the genotype of the mice in all experiments and analyses.

### Neurovascular Coupling and CO_2_ stimulation - Laser speckle imaging

The mice were anaesthetised with ketamine (70 μg/g body weight, bela-pharm) and xylazine (16 μg/g body weight, Bayer AG). The body temperature was maintained at 37±0.2°C throughout the study using a feedback-regulated warming system (TCAT-2LV, Physitemp Instruments). A small ventilatory tube was inserted into the trachea after tracheotomy was performed and connected to a small animal ventilation device (MiniVent, Harvard Apparatus). The mice were ventilated with a gas mixture reflecting normal air (21% O_2_, 79% N_2_). The ventilation volume was constant, and the ventilation frequency was adjusted to a physiological expiratory CO_2_ concentration of 35-45 mmHg that was continuously controlled during the experiments with a capnometer (microCapstar, CWE Inc.). The skin over the head was removed, but the skull remained intact for the whole experiment. Laser speckle imaging (FLPI, Moor instruments) was carried out using a recording rate of 0.25 Hz during CO_2_ stimulation or 25 Hz to measure neurovascular coupling responses. Regions of interest were defined over the somatosensory cortex (see Fig. 1D), and the flux intensities were normalised to the baseline values. CO_2_ stimulation was performed using a gas mixture containing 10% CO_2_, combined with 21% O_2_ and 69% N_2_. To measure neurovascular coupling, the whisker pad was electrically stimulated for five seconds using bipolar pulses (750 µA, 1 ms, 5 Hz; STG4002, Multichannel systems) after inserting two needle electrodes subcutaneously as described before.^23^ Stimulation was performed five times, and the mean curves of each mouse were analysed.

**Figure 1:**
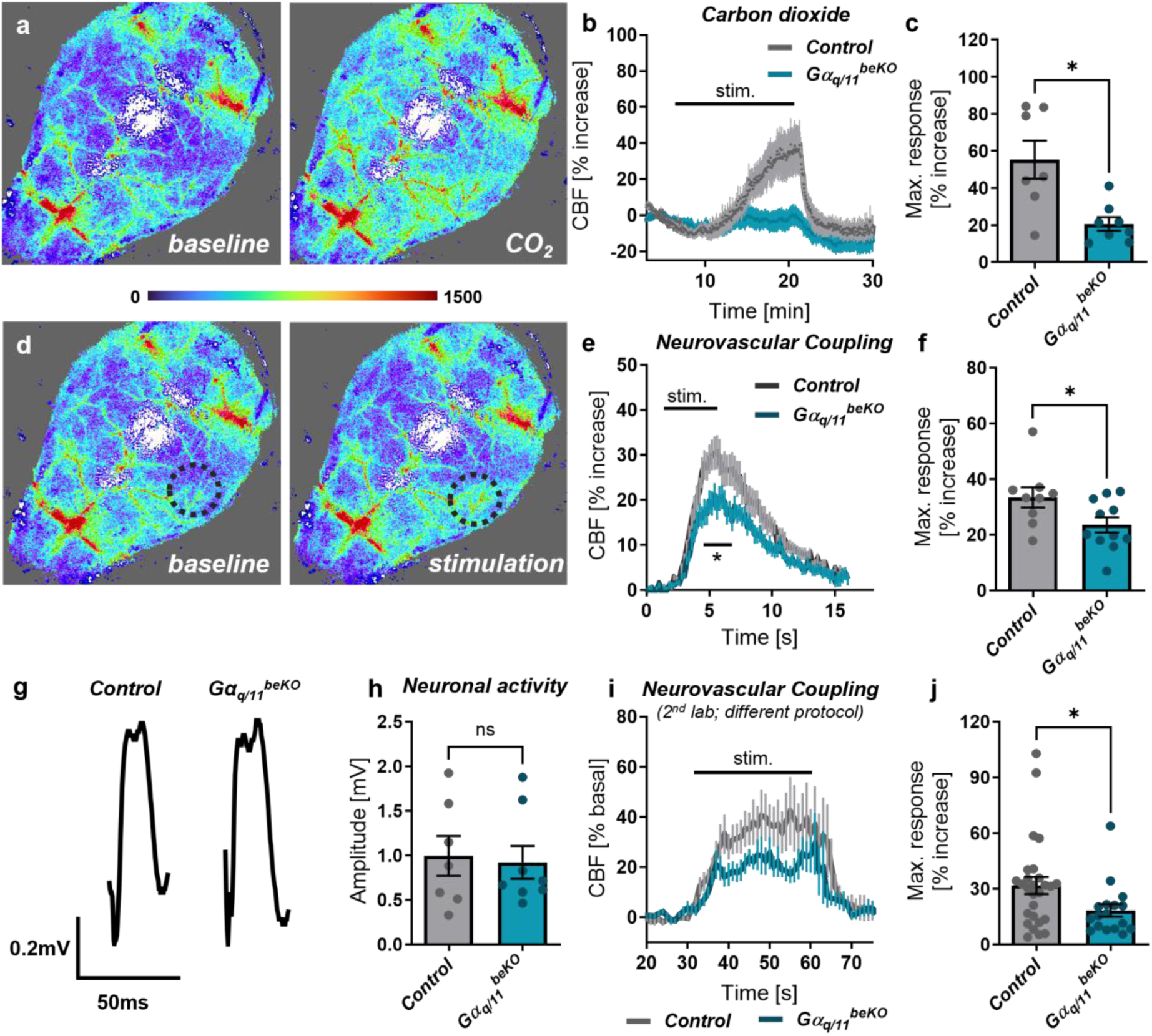
***Brain endothelial Gα_q/11_ signalling mediates cerebrovascular reactivity to CO_2_ and neuronal activity.*** (**a**) Representative images of perfusion measurements using laser speckle imaging before and during CO_2_ exposure. (**b**) Time course of cerebral blood flow (CBF) during CO_2_ exposure in ventilated, ketamine/xylazine-anaesthetised Gα_q/11_^beKO^ (mice carrying a brain endothelial-specific deletion of Gα_q/11_) and control mice. N=7-8. (**c**) Maximal CBF response of curves shown in (b). (**d**) Representative images of perfusion measurements using laser speckle imaging before and during whisker pad stimulation. Black circle indicates the analysed area above the somatosensory cortex. (**e**) Time course of cerebral blood flow (CBF) during whisker pad stimulation in ventilated, ketamine/xylazine-anaesthetised Gα_q/11_^beKO^ and control mice. N=9-11. (**f**) Maximal CBF response of curves shown in (e). (**g**) Representative traces of local field potentials measured in Gα_q/11_^beKO^ and control mice during whisker pad stimulation. (**h**) Mean amplitudes of local field potentials measured in Gα_q/11_^beKO^ and control mice. (**i**) Time course of CBF measured by laser Doppler flowmetry during whisker pad stimulation in spontaneously breathing, isoflurane-anaesthetised Gα_q/11_^beKO^ and control mice. N=17-27 recordings of 6-9 mice. (**j**) Maximal CBF response of curves shown in (i). Shown are means±SEM. ns: p>0.05, *p<0.05. **p<0.01. Detailed information on the age and time after tamoxifen, as well as the exact test statistics and values, is provided in Supplemental Table 3.

### Neurovascular Coupling - laser Doppler recording

The mice were anaesthetised with 1.5-2% isoflurane in a mixture of 40% O_2_ and 60% N_2_, and were allowed to breathe spontaneously through a head mask. The body temperature was maintained at 37°C with a servo-controlled heating pad. The animal was fixed in a stereotaxic frame, and a craniotomy was performed in the right parietal bone using a drill. The dura in the cranial window was carefully dissected, and the tissue was kept wet with aCSF (pH 7.4). A laser Doppler needle probe (Periflux 5000, Perimed AB) was positioned at the somatosensory cortex. Neurovascular stimulation was performed as previously described.^24^ In brief, whiskers were stimulated electromechanically with a bending actuator (#PL112-PL140; PICMA) connected to a piezo amplifier (#E-650 Amplifier, Physik Instrumente) according to the following protocol: the left whiskers were stimulated for 30 s, repeated 3 times, 60 s apart. Stimulation-evoked perfusion responses in the contralateral cortex were recorded using the laser Doppler probe.

### Local field potential (LFP) recordings

LFPs were measured as described before.^25^ Briefly, mice were prepared in a similar way to that for laser speckle imaging, including anaesthesia with ketamine/xylazine and artificial ventilation (see above). Two holes were drilled to place the electrodes (Tungsten microelectrodes; FHC), the first one over the cerebellum for the reference electrode and a second one over the barrel cortex for the recording electrode. The whisker pad was stimulated in a similar way to the neurovascular coupling stimulation described above, using electrodes inserted into the whisker pad and an electrical stimulator. The stimulus was the same as described before, too (see above). LFPs were recorded with an amplifier (Model 1800; AM-Systems) and converted into a digital format by an AD converter (PowerLab 8/30 ML870; ADInstruments). LabChart 7 (ADInstruments) was used to record and analyse the data (sampling rate 20 kHz; notch filter 50 Hz). For each mouse, four to six different areas in the contralateral barrel cortex were recorded to find the region with the highest response upon whisker stimulation. Here, each region was stimulated three times. Then the amplitudes of 30 pulses taken from the region having the highest amplitude were averaged for each mouse.

### Arterial Spin Labelling MRI measurements and Time-of-Flight (TOF) MR angiography

Anaesthesia was induced by 4% and maintained at 0.8-1% isoflurane in 21% O_2_ and 79% N_2_. The respiration rate and body temperature were continuously monitored using an MRI-compatible system, and the temperature was maintained at 37±0.2°C throughout the study using a feedback-regulated warming system (Small Animal Instruments Inc.). The mice were scanned using a 7 Tesla small animal MRI scanner (ClinScan, Bruker). For analyses of the cerebral arterial vasculature, the TOF data sets were analysed using IMARIS (v9.9.1, Bitplane) as illustrated in Supplemental Fig. 5. For detailed settings and analyses, see Supplemental Methods.

### Two-photon microscopy

Mice were anaesthetised with urethane (1.8-2.0 mg/g body weight), tracheotomized, and artificially ventilated (79% N_2_, 21% O_2_; MiniVent, Hugo Sachs Elektronik). The ventilation volume was constant, and the ventilation frequency was adjusted to a physiological expiratory CO_2_ concentration of 35-45 mmHg that was continuously controlled during the experiments with a capnometer (microCapstar, CWE Inc.). Tetramethylrhodamine B thiocarbamoyl (TRITC) dextran (2%, 150 kDa, TdB consultancy) was injected intravenously to visualise the vasculature. In a stereotactic frame, an open cranial window was drilled and covered with an artificial tears gel (Bausch&Lomb) to bridge the working distance of the water immersion objective used. Images were acquired with a two-photon microscope (TRiM Scope I, LaVision Biotec), equipped with an optical parametric oscillator Titan-Saphir laser combination (Coherent Inc.). Image stacks of 1×1×0.5 mm volume with a step size of 2 µm were scanned during the excitation of TRITC with a wavelength of 1100 nm. Line scans along single vessels were performed with a scanning frequency of 800-900 Hz for one second to determine the flow speed of red blood cells and the direction of blood flow. To assess arteriolar and venular vessels, at least three branches of the same vascular tree were scanned to define converging or diverging vessels. ImageJ software was then used to analyse vessel diameters and blood cell speed after stacking the line scans.

### Object place recognition test

The object place recognition (OPR) test was performed with appropriate adaptations as described earlier.^26^ First, mice underwent a habituation phase once per day for three days, during which they were placed in the centre of a dimly lit (approximately 20 lux) rectangular open field apparatus and left for 10 min to freely explore it. The next day, the sample trial and the test trial took place. In the sample trial, two identical objects were placed in the corners of the apparatus, and the mice were allowed 10 min to explore the apparatus and the objects freely. Afterwards, the mice were transported back to their home cage for 60 min. During the test trial, the mice were placed in the open field apparatus and allowed for five minutes to explore the previously presented objects, with the difference being that one object was displaced. A visual clue attached to the wall of the apparatus was used for the spatial orientation of the mice.^27^ The animals’ behaviour during the OPR test was recorded with a digital camera, and the scoring procedure was conducted using the software ANY-maze (Stoelting Inc.). The preference index was calculated as the ratio of time spent exploring the displaced object divided by the total time for exploring both objects. Different combinations of objects and objects’ locations were used between animals, and for each time point, to eliminate any possible biased preferences.

### Barnes maze test

The Barnes maze test was performed as described previously^28^ with minor adaptations. The Barnes maze apparatus consisted of an elevated circular platform with 19 closed holes and one open hole leading to an escape box (Stoelting Co., Cat. Nr. 60170). The mice were trained to find the escape box based on visual cues placed in the surrounding environment. The apparatus was brightly lit with approximately 350 lux. The mice underwent five days of acquisition trials followed by the probe trials on days 6 and 17. During the acquisition trials, the mice were placed in the middle of the apparatus and allowed to find the escape box for three minutes. If the mice failed to locate or enter the escape hole in this period, they were gently guided and placed inside the escape box. The mice were left inside the escape box for 1 min before being transported back to the home cage. Each mouse received four acquisition trials daily with an inter-trial interval of 15 min. On days 6 and 17, the escape box was removed, the hole was closed, and the apparatus was turned by 180° to exclude the influence of intra-maze cues. Then, mice were placed in the middle of the apparatus and allowed to explore it for 90 s before being transported back to their home cage. The mice were recorded during acquisition and probe trials, and the videos were analysed offline using ANY-maze (Stoelting Inc.). Primary latency, primary distance, and primary errors were defined as time, distance, and the number of pokes into other holes before finding the target hole, respectively.

### Tissue processing for subsequent protein analyses

Mice were transcardially perfused with heparin (2 or 10 IU/mL)/Ringeŕs solution under deep anaesthesia. Brains were removed and either snap-frozen directly or post-fixed in 3.7% formalin solution at 4 °C overnight. After post-fixation, the brains were washed and left for 48 h in a 30% saccharose solution. Finally, the brains were snap-frozen in 2-methylbutane and kept at −80 °C until cryosectioning was performed.

### Immunohistochemical staining and microscopy

Brains were cut with a cryotome (CM3050, Leica) at −25 °C at a thickness of 20 µm and stored at −20 °C. For staining, the sections were permeabilised with 0.3% Triton-X 100 for 15-30 min and then blocked with 3-5% BSA in 0.3% Triton-X 100 for 30-75 min at room temperature. Then, sections were incubated with the respective primary antibody diluted in the blocking solution at 4 °C overnight. Incubation with secondary antibodies took place for one hour at room temperature. For collagen IV, caveolin-1, shank2, MAP2, and bassoon staining, an antigen retrieval step (20 min boiling at 95 °C in a 10 mM tri-sodium citrate buffer pH 6.0) was included. For the MAP2 staining, a Fab fragment solution (115-007-003, Jackson ImmunoResearch) was added to the blocking solution to reduce the nonspecific binding of the primary antibody. All the antibodies used are listed in Supplemental Tables 1 and 2. The images were acquired using the fluorescence microscopes Leica DMI6000B and Leica Stellaris 5.

### Image analyses

Analyses were performed in a blinded manner using ImageJ (National Institutes of Health). Multi-channel microscopy images were batch-processed using custom Fiji ImageJ (National Institutes of Health, Version 1.54r) macros to segment markers of interest via automated thresholding and morphological filtering. Selection of suitable thresholds and pre-processing steps was performed in a blinded manner. The resulting binary masks were used to calculate mean signal intensities, vessel area coverage, and marker spatial overlap within specific compartments. Furthermore, masks of vessel markers (e.g., collagen IV, CD31) were skeletonised to quantify total vessel length.

String vessels were analysed manually and identified as structures positively stained for the basement membrane component collagen IV and negatively stained for an endothelial marker, as described earlier.^29^

### Immunoblotting

Hippocampal and cortical tissue samples were prepared by cutting the respective brain regions from frozen brains in the cryotome. Primary brain endothelial cells (see PBECs section) were snap-frozen in liquid nitrogen after treatment. Tissue or PBECs were lysed, and the protein content was determined using a Lowry assay. Samples were supplemented 1:4 with SDS buffer (0.75 M Tris-HCl, 0.08 g/mL SDS, 40% glycerol, 0.4 mg/mL bromophenol blue, and 62 mg/mL DTT) and incubated at 95 °C for ten minutes. After loading on SDS–PAGE gels, proteins were transferred to nitrocellulose membranes, which were then incubated with primary antibodies overnight at 4 °C. Afterwards, the membranes were incubated with HRP-conjugated secondary antibodies for two hours at room temperature. For detection, enhanced chemiluminescence (SuperSignal West Femto Substrate, Thermo Scientific) and a digital detection system (Fusion Solo S, Vilber) were used. Immunoblots were analysed by the detection software or ImageJ (National Institutes of Health. All the antibodies used are listed in Supplemental Tables 1 and 2.

### Preparation or primary brain endothelial cells (PBECs)

PBECs were prepared from adult wild-type or *Cdh5*-GCaMP8 mice, or adult Gα_q/11_^beKO^ and control mice two weeks after tamoxifen injections. Cells were prepared as described previously^30^ and used for experiments when reaching a confluency of at least 80%. For VEGF (50 ng/mL, mVEGF-164. R&D systems) or carbachol (100 µM, Sigma-Aldrich) stimulation, cells were placed in a physiological salt solution (NaCl 130 mM, NaH_2_PO_4_ 1.25 mM, KCl 3 mM, NaHCO_3_ 26.5 mM, MgCl_2_ 2 mM, CaCl_2_ 2 mM, D-glucose 10 mM) before stimulation that was pregassed with 5% CO_2_ at 37 °C. For blocking the Gα_q/11_ signalling pathway, PBECs were pre-incubated with 10 μM of the specific inhibitor FR900359^31^ or 0.1% DMSO as vehicle control for 30 min at 37 °C in physiological salt solution before measurements. Supernatant of PBECs was collected for NO release assay (see below) after 30 min at 37 °C in physiological salt solution and frozen in liquid nitrogen for further analyses.

### Nitrate asssay

NO release in the supernatant of PBECs was determined according to the manufactureŕs protocol using a nitrate/nitrite fluorometric assay kit (CAY780051-192, Cayman Chemical).

### Calcium imaging

Imaging of intracellular Ca^2+^ changes was performed in two different ways. For the detection of stimulation-induced calcium increases, we made use of PBECs from *Cdh5*-GCaMP8 mice. ^21,22^ PBECs were prepared as mentioned, and fluorescence was measured at a fluorescence microscope during application of VEGF (50 ng/mL) or carbachol (100 µM).

To measure spontaneous Ca^2+^ activity during a 30-min time course, PBECs were cultured on cover glasses and stained as described previously ^32^. Briefly, PBECs were incubated with Fluo-4 AM (5 μM with 0.05% pluronic 127, and 10 μM probenecid, all from Life Technologies) in physiological salt solution with or without 10 μM of the specific Gα_q/11_ inhibitor FR900359 (see PBEC preparation) for 30 min at 37 °C. Then, they were incubated in salt solution containing only probenecid (10 μM, 30 min 37 °C) before measurement. PBECs were placed in a flow chamber that was perfused (flow rate: 2 ml/min) and continuously gassed with 5% CO_2_ / 95% O_2_. Fluorescence was measured using a high-speed calcium imaging setup (FEI GmbH). Changes in the fluorescence ratio were normalised to baseline mean values during the first 30 sec (F_0_), and changes are shown as F/F_0_. Several cells per sample were randomly chosen and analysed using ImageJ. For assessing spontaneous calcium signals, one calcium event was defined as a reversible increase of at least 5 % in F/F_0_. The data sets were analysed for the percentage of responding cells with at least one event during a 30-min recording frequency of Ca^2+^ signals (# of events/hour).

### Statistics

No statistical methods were used to predetermine sample sizes, but our sample sizes are similar to those reported in previous publications.^15,25,29^ All the values are expressed as means ± SEM. Analyses were performed using GraphPad Prism 10.4. Data values were assessed for normality by the Shapiro-Wilk or the Anderson-Darling test, for equal variances using the F test, and for homoscedasticity by Spearmańs test. If the raw data did not meet these criteria, a non-parametric method was applied. Two groups were compared using a two-tailed, unpaired, parametric Student’s t-test or Mann-Whitney test. For multiple comparisons, we used one- or two-way ANOVA or Scheirer-Ray-Hare tests, if data set was not suited for ANVA analyses. A Sidak’s post-hoc or Mann-Whitney post-hoc test with Bonferroni-Sidak correction was performed if the ANOVA or Scheirer-Ray-Hare test showed a significant difference. Survival was tested using Kaplan-Meier methods, and curve comparison was performed using the Log-rank (Mantel-Cox) test. Differences were considered to be significant at p < 0.05. If more than one test was applied (e.g., post-hoc tests), we corrected for multiple comparisons and reported adjusted p values. All statistical details of each experiment are listed in Supplemental Table 3. The investigators were blinded to the different groups in all experiments and analyses.

## RESULTS

### Brain endothelial Gα_q/11_ signalling mediates cerebrovascular reactivity

In order to ascertain the function of the endothelial Gα_q/11_ pathway in cerebrovascular vasodilation, we exposed mice lacking both Gα_11_ and Gα_q_ specifically in the brain endothelium (Gα_q/11_^beKO^) to carbon dioxide (CO_2_). As previously demonstrated,^15^ the acute responsiveness to CO_2_ was significantly reduced (Fig. 1a-c). Additionally, Gα_q/11_^beKO^ mice exhibited an impaired vascular response to neuronal activity induced by whisker pad stimulation (Fig. 1d-f). In comparable experiments, we did not observe any difference in neuronal activity as measured by local field potentials (Fig. 1g, h), thus confirming an acute endothelium-mediated mechanism acting downstream of neuronal activity.

The genetic strategy used to induce the deletion of the Gα_q/11_ signalling pathway in brain endothelial cells involves the whole-genome deletion of *Gna11* and the inducible deletion of *Gnaq* using the brain endothelial-specific driver mouse line *Slco1c1*-CreER^T2^.^16^ We confirmed the successful deletion by qPCR and Western blotting using primary brain endothelial cells (PBECs) of these mice (Supplemental Figure 1a, b). In *Gna11* knockout mice, the neurovascular coupling response was similar to that in control mice (Supplemental Fig. 2a, b), indicating that whole-genome deletion of Gα_11_ alone does not interfere with neurovascular coupling. Additionally, as *Slco1c1*-CreER^T2^ mice show some minor recombination in astrocytes,^16^ we also assessed the neurovascular coupling response in astrocyte-specific Gα_q/11_ knockout mice using a Glast-CreER^T2^ mouse line (Gα_q/11_^astroKO^).^18^ In these mice, the perfusion response to whisker pad stimulation was similar to that in control mice (Supplemental Fig. 2c, d), supporting the finding that the early phase of vasodilatory responses to neuronal stimulation is not mediated by intracellular calcium release via Gα_q/11_-IP_3_R signalling in astrocytes.^33^ However, prolonged stimulation might have produced different results since astrocytic calcium is involved in the later, sustained phase of neurovascular coupling.^34^ To confirm the specific brain endothelial involvement in cerebrovascular reactivity, we additionally applied an alternative viral strategy based on a brain endothelial cell-specific AAV vector.^20^ By using the AAV-BR1 capsid, we transduced specifically the brain endothelium and expressed a Cre recombinase in *Gna11*^-/-^; *Gnaq*^fl/fl^ mice (Gα_q/11_^BR1-beKO^). Similar to the *Slco1c1*-CreER^T2^ approach, we achieved a successful reduction of *Gnaq* mRNA expression, measured in primary brain endothelial cells from Gα_q/11_^BR1-beKO^ and control mice (Supplemental Figure 3a). Also, in those mice, the loss of the Gα_q/11_ signalling pathway in brain endothelial cells led to a reduced response to the vasodilating stimulus CO_2_ (Supplemental Fig. 3b, c).

Brain endothelial Gα_q/11_ proteins have been shown to affect ion channel activities, some of which affect neurovascular coupling responses.^11,22,35^ Nevertheless, it is still not clear whether endothelial Gα_q/11_ proteins themselves are involved in the vasodilation upon neuronal activity. To validate our findings, we repeated the experiments in a different laboratory environment using alternative anaesthesia and without artificially ventilating the mice. Consistent with the former results, we found that Gα ^beKO^ mice exhibited a reduced perfusion response to neuronal activity compared to control mice (Fig. 1i, j). In summary, the vascular response to neuronal activity or CO_2_ is strongly impaired in Gα_q/11_^beKO^ mice, indicating a cerebral endothelial dysfunction similar to the human situation in cSVD.^2,4^

*Gα_q/11_^beKO^ mice do not show structural or functional vascular alterations at baseline* Functional impairment of the braińs vasculature may be accompanied or even caused by morphological alterations of the vessel structure. Previously, we demonstrated that the endothelial cells of the main arteries in the brain show recombination in *Slco1c1*-CreER^T2^ mice.^25^ Tissue sections from the head-supplying arteries using the reporter mouse line *Slco1c1*-CreER^T2^; Ai14^+/fl^ ^19^ confirmed that the recombination leading to tdTomato expression in this mouse line affected the internal carotid artery starting from the bifurcation but not the external carotid artery (Supplemental Fig. 4). We next sought to determine whether changes in brain-supplying vessels brought about functional impairment in Gα_q/11_^beKO^ mice and used two techniques – time-of-flight (TOF) angiography and two-photon microscopy – to trace the vasculature from the arterial level down to the capillary level in living mice.

The three-dimensional reconstruction of TOF angiography (Fig. 2a, Supplemental Fig. 5) and analyses of two main arterial vascular trees, namely the middle and the posterior cerebral artery (MCA and PCA, respectively), revealed no major differences in terms of diameter, overall volume, or straightness between control and Gα_q/11_^beKO^ mice (Fig. 2a-h), although the PCA and its branches in Gα_q/11_^beKO^ mice showed a little reduction in straightness and had slightly higher volumes than those in control mice (Fig. 2f, h). However, this technique has a limited spatial resolution and only detects large arteries and arterioles on the surface of the brain. As vascular resistance and flow regulation are primarily mediated by smaller arterioles and first-order capillaries,^36^ we used two-photon microscopy to examine the vasculature in greater detail (Fig. 2i). We recorded volumes of the cortical vasculature along entire vascular trees (Fig. 2i, left panel) as well as fast line scans along individual vessels to calculate blood flow velocity (Fig. 2i, right panels). No differences were found between control and Gα_q/11_^beKO^ mice in terms of arteriolar or capillary diameter (Fig. 2j, l) or red blood cell speed (Fig. 2m), thus extending the

**Figure 2:**
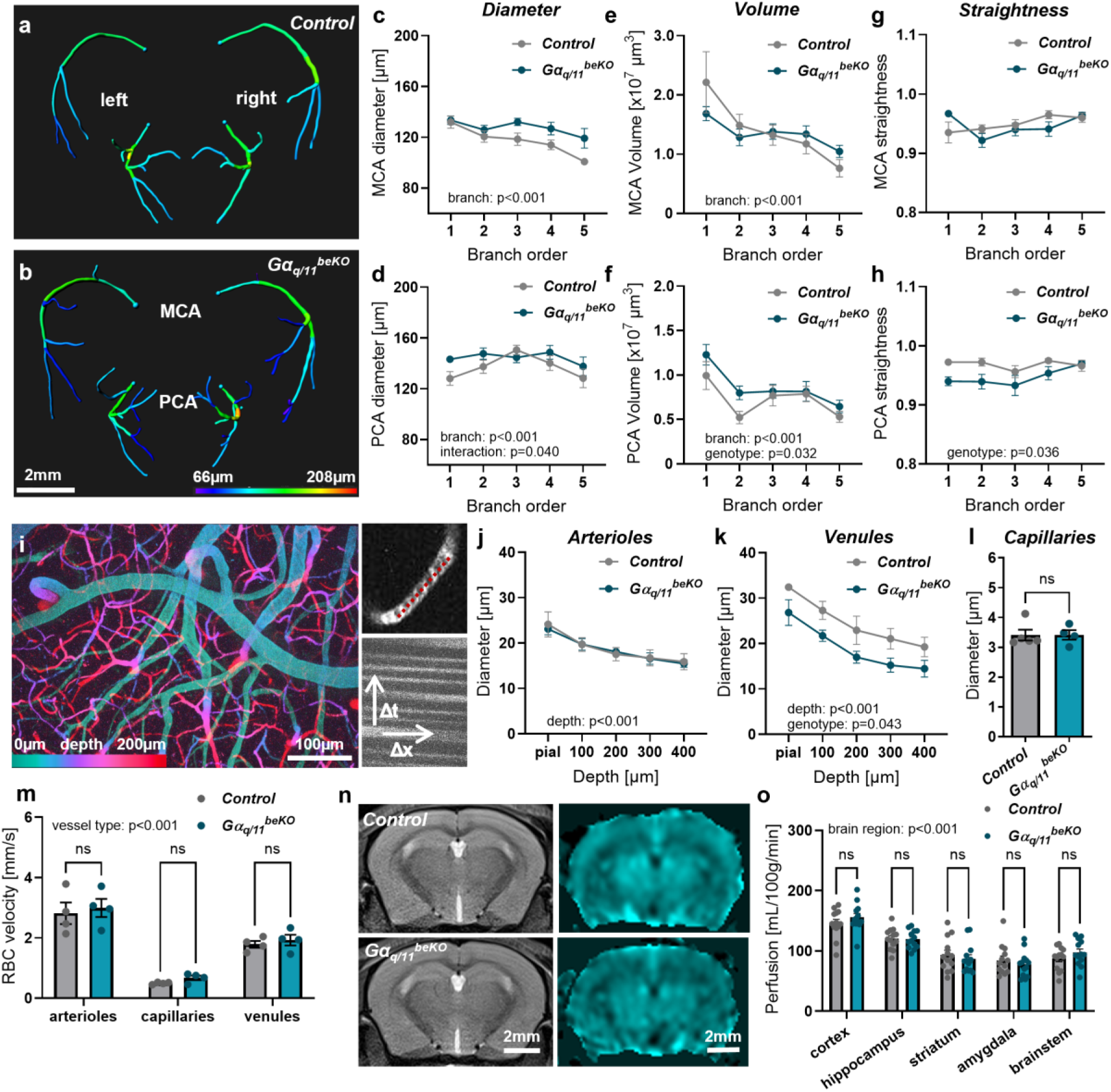
***Gα_q/11_^beKO^ mice do not show structural or functional vascular alterations at baseline.*** (**a**, **b**) Representative images of reconstructed middle and posterior cerebral artery branches (MCA, PCA) from time-of-flight (TOF) angiography measurements in control (a) and Gα_q/11_^beKO^ (b) mice. Colour-coding indicates diameter of the respective vessel segment. See Supplemental Figure 5 for detailed reconstruction processing. (**c-h**) Quantifications of diameter, volume, and straightness of vascular branches of MCAs and PCAs of control and Gα_q/11_^beKO^ mice. Branch order 1 refers to the first segment originating from the Circle of Willis. Means of left and right arteries are shown. N=8-14. Significantly different variables are indicated in the graphs. (**i**) Maximum intensity projection of a volume taken by two-photon microscopy is shown (left side). Tissue depth is colour-coded. Right upper panel: a single capillary filled with blood cells (dark gaps) is shown. Red line indicates a line scan to measure red blood cell (RBC) velocity. Stacking of line scans enables determination of velocity by taking the length of the line and the frequency of the scanning into account (right lower panel). (**j-l**) Diameter of penetrating arterioles (j), ascending venules (k), and capillaries (l) of control and Gα_q/11_^beKO^ mice are shown. N=4-5. Significantly different variables are indicated in the graphs. (**m**) Quantification of RBC velocities in different parts of the vascular tree of control and Gα_q/11_^beKO^ mice is shown. N=4. Significantly different variable is indicated in the graph. (**n**) Representative images of T2-weighted magnetic resonance imaging (MRI) and arterial spin labelling (ASL) MRI of control and Gα_q/11_^beKO^ mice are shown. T2 images were used to define regions to be analysed in the ASL images. (**o**) Quantification of ASL signals in different brain regions of control and Gα_q/11_^beKO^ mice is shown. N=13. Significantly different variable is indicated in the graph. Shown are means±SEM. ns: p>0.05. Detailed information on the age and time after tamoxifen, as well as the exact test statistics and values, is provided in Supplemental Table 3.

TOF-angiography results to smaller blood-supplying vessels. Interestingly, the diameter of venules was smaller in Gα ^beKO^ compared to control mice (Fig. 2k) without affecting the RBC velocity (Fig. 2m), suggesting an impairment of the passive venular part of the vascular tree and might hint at an impaired blood drainage. Finally, we used arterial spin labelling MRI to determine whether these small vascular alterations affected overall perfusion in several brain regions; however, we found no differences between Gα ^beKO^ and control mice (Fig. 2n, o).

*Brain endothelial Gα_q/11_ signalling affects calcium, NO, and phosphorylation of tau* Endothelial dysfunction is a feature of cSVD, which is linked to cognitive impairment.^37–39^ It has been recently shown that cerebral endothelial dysfunction induced by high dietary salt gives rise to increased tau phosphorylation due to reduced nitric oxide (NO) release, leading to cognitive impairment.^40^ Hyperphosphorylation of tau is a hallmark of certain neurodegenerative diseases and causes cognitive impairment via neuronal dysfunction.^41^ NO release is strongly induced by the activation of endothelial Gα_q/11_ signalling, which increases intracellular calcium concentration and activates the endothelial NO synthase, and this could provide a link between cerebrovascular and cognitive dysfunction in Gα ^beKO^ mice. When inhibiting the Gα_q/11_ pathway by using the specific inhibitor FR900359^31^ in primary brain endothelial cells, we could indeed see that spontaneous calcium events were significantly diminished (Fig. 3a-c), accompanied by a reduced NO release from these cells (Fig. 3d). Successful inhibition of the Gα_q/11_-mediated coupling between receptor and increased calcium was demonstrated by stimulating primary brain endothelial cells of *Cdh5*-GCaMP8 mice^21^ with carbachol, a muscarinic acetylcholine receptor agonist (Fig. 3e-g). Since spontaneous events as well as stimulated intracellular calcium increase in brain endothelial cells occur in the brain during neuronal activity,^22^ we hypothesised that NO-mediated regulation of tau could be altered in Gα ^beKO^ mice. Therefore, we measured tau phosphorylation in the hippocampus and cortex of Gα_q/11_^beKO^ and control mice. Indeed, we found increased phosphorylation (Fig. 3h-k) at epitopes that promote tau aggregation and neuronal dysfunction in Gα ^beKO^ mice (pSer202/pThr205 by AT8 and pThr231 by AT180 antibodies).^41^ In summary, the loss of Gα_q/11_ signalling in brain endothelial cells induces an isolated functional impairment of the brain vasculature, leading to an increased phosphorylation of tau. Hence, we investigated the cognitive consequences of that cerebral endothelial dysfunction.

**Figure 3:**
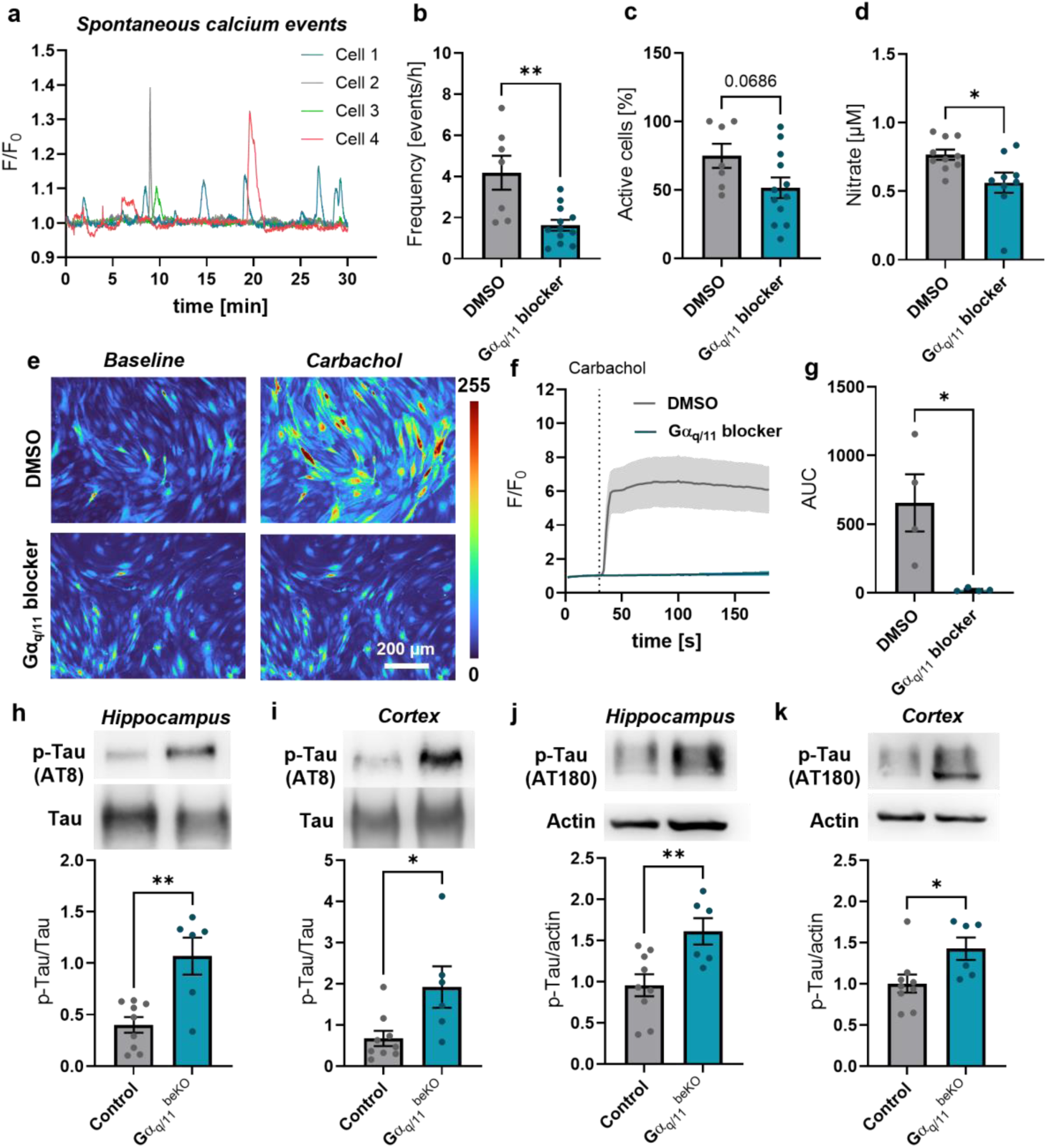
***Brain endothelial Gα_q/11_ signalling affects calcium, NO, and phosphorylation of tau.*** (**a**) Representative spontaneous calcium events measured in primary brain endothelial cells over 30 min. (**b, c**) Quantification of frequency (b) of spontaneous calcium events and percentage of active cells (c) measured in primary brain endothelial cells stained with Fluo-4 and pre-treated with DMSO (vehicle) or the Gα_q/11_ blocker FR900359 (10 µM). N=7-12. (**d**) Nitrate concentration measured in the supernatant of primary brain endothelial cells pre-treated with DMSO (vehicle) or the Gα_q/11_ blocker FR900359 (10 µM). Incubation period was 30 min. (**e-g**) Representative images (e) and quantification (f, g) of the calcium response in primary brain endothelial cells from *Cdh5*-GCaMP8 mice to the muscarinergic acetylcholine receptor agonist carbachol (100 µM) pre-treated with DMSO (vehicle) or the Gα_q/11_ blocker FR900359 (10 µM). N=4. (**h-k**) Representative Western blots and quantification of phosphorylated (h, i: AT8, pSer202/pThr205; j, k: AT180, pThr231) and full tau (h, i) or actin (j, k) in hippocampal (h, j) and cortical (j, k) tissue of control and Gα_q/11_^beKO^ mice are shown. N=6-9. Shown are means±SEM. *p<0.05, **p<0.01. Detailed information on the age and time after tamoxifen, as well as the exact test statistics and values, is provided in Supplemental Table 3.

### Loss of brain endothelial Gα_q/11_ signalling leads to mild cognitive impairment in aged mice

To test memory, we performed object-place recognition tests (Fig. 4a) before inducing the knockout at eight weeks of age (Fig. 4b) and at various time points subsequently during the lifespan of the same mice (Fig. 4c-e). Interestingly, only Gα ^beKO^ mice at the age of 18 months failed to reach a significant preference for the replaced object, indicating a mild cognitive deficit (Fig. 4e) without showing a significant impairment in general motor activity (Supplemental Fig. 6a, b). To further investigate memory function in these old mice, we performed a Barnes maze test when they were 20-22 months old (Fig. 4f). During the learning phase of the test, control mice displayed the expected improvement in their search strategy with the most of the single tests being solved by directly finding the hole on day five (Fig. 4g, i). Gα_q/11_^beKO^ mice, in contrast, had a marked improvement from day one to day two of the learning phase but failed to improve further in subsequent days (Fig. 4h). On day five, they utilised significantly fewer direct strategies to find the hole than the control mice did (Fig. 4i). Also, the primary latency, until the hole was found, was higher in Gα_q/11_^beKO^ mice throughout almost the whole learning phase (Fig. 4j). Lastly, on the test days, Gα_q/11_^beKO^ mice performed worse than control mice, showing a significantly higher number of primary errors and latency on day six, but not on day 17 (Fig. 4k, l), indicating mild cognitive deficits and prolonged duration of memory retrieval. Although the total distance travelled during the tests (Supplemental Fig. 6c) was lower in Gα ^beKO^ mice, the primary distance was not different in Gα ^beKO^ compared to control mice (Supplemental Fig. 6d), arguing against an influence of a different motor activity on memory test outcome.

**Figure 4:**
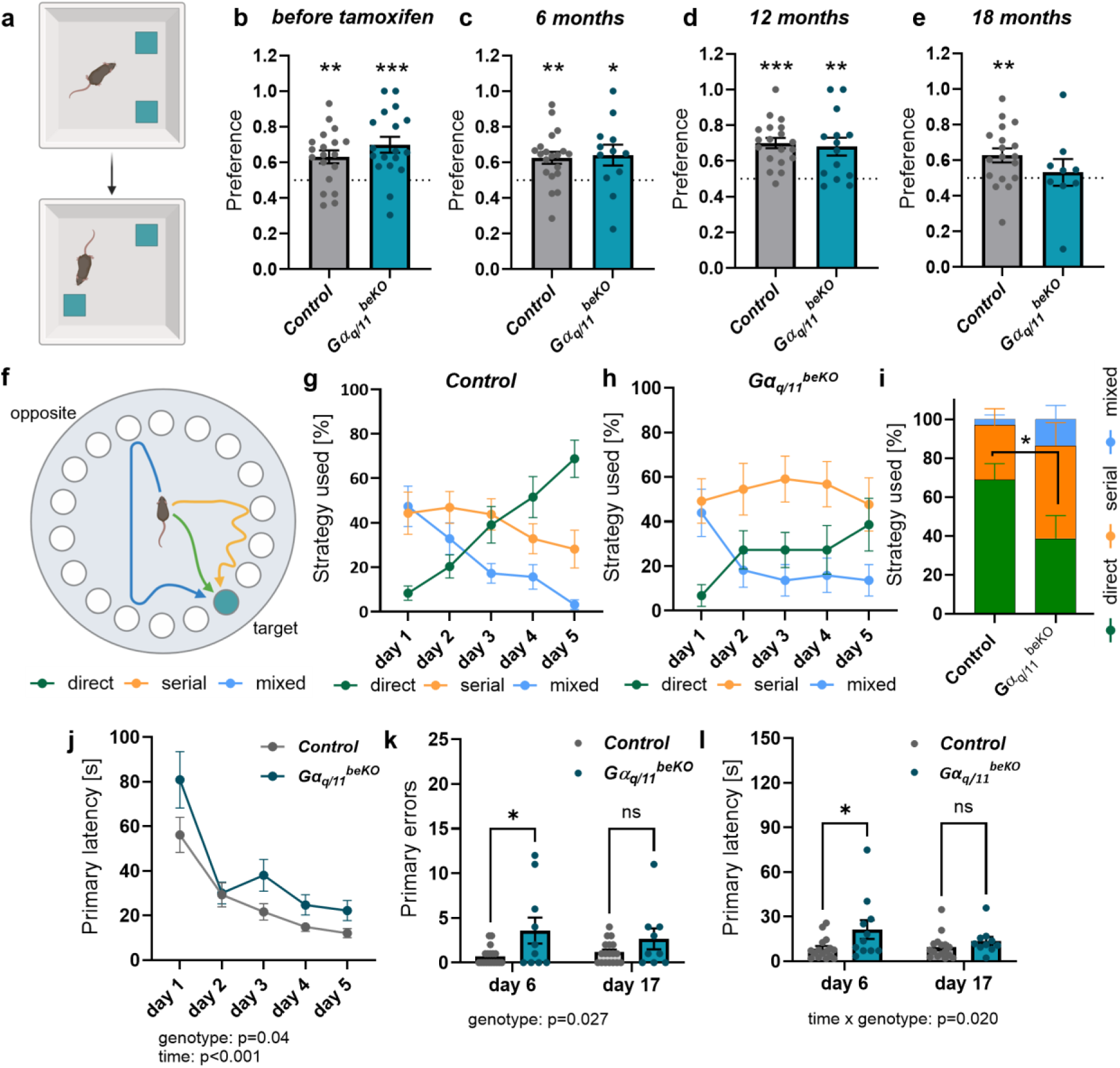
***Loss of brain endothelial Gα_q/11_ signalling leads to mild cognitive impairment in aged mice.*** (**a**) Scheme of object place recognition (OPR) test with the upper part showing the situation before and the lower part showing the situation after moving one of the two objects. (**b-e**) OPR preference indices at different time points are shown. Tamoxifen was administered at the age of 7-8 weeks to control and Gα_q/11_^beKO^ mice to induce genetic deletion. Time points indicate the age of the mice. Dotted lines mark chance level (0.5), and Student’s *t*-tests were applied to test against chance level. N=9-18. (**f**) Scheme of Barnes maze test is shown with different colours indicating different strategies to find the target hole. (**g**, **f**) The proportion of different strategies on the training days is shown for control (g, N=16) and Gα_q/11_^beKO^ (h, N=11) mice. (**i**) Comparison of strategies used on day 5 of the training between control and Gα_q/11_^beKO^ mice. N=11-16. (**j**) Primary latency until the target hole was found during the training days is shown for control and Gα_q/11_^beKO^ mice. N=11-16. Significantly different variables are indicated in the graph. (**k**, **l**) Primary errors (k) and primary latency (l) on test days 6 and 17 are shown for control and Gα_q/11_^beKO^ mice. N=11-16. Significantly different variables are indicated in the graph. Shown are means±SEM. ns: p>0.05, *p<0.05, **p<0.01, ***p<0.001. Detailed information on the age and time after tamoxifen, as well as the exact test statistics and values, is provided in Supplemental Table 3.

### Aged Gα_q/11_^beKO^ mice show structural alterations in the white matter of the brain

To investigate the underlying alterations in the brains of aged Gα_q/11_^beKO^ mice, we performed immunofluorescence staining of neuronal and synaptic markers showing that the number of neurons and synapses in the hippocampus and cortex of aged Gα_q/11_^beKO^ mice was unchanged (Supplemental Fig. 7 and Supplemental Fig. 8a, b), indicating normal cellular morphology and a functional deficit in these brain regions rather than a morphological one. In addition, the expression of brain-derived neurotrophic factor (BDNF), a neuroprotective protein^42^ whose presence in endothelial cells is essential for memory function,^43^ was not altered in cortical and hippocampal tissue of aged Gα_q/11_^beKO^ mice as well (Supplemental Fig. 8c, d). However, early markers of vascular dementia in humans include white matter alterations with increased hyperintensities indicating disturbed myelination.^2,39^ Therefore, we examined the myelination levels by staining for myelin basic protein (MBP) in grey and white matter areas in different regions along the mouse brain (Fig. 5a-d). We found decreased myelination in the posterior part of the forebrain, which was most prominent in the white matter of the corpus callosum (Fig. 5c, d). Together with increased tau phosphorylation, this finding suggests that brain endothelial Gα_q/11_ signalling protects against neuronal dysfunction in aged mice. Interestingly, although microglial activation is reported in cSVD,^44^ we observed no signs of neuroinflammation in different brain regions of aged Gα_q/11_^beKO^ mice, as indicated by staining for microglial and astrocytic activation markers (Supplemental Fig. 8e-i).

**Figure 5:**
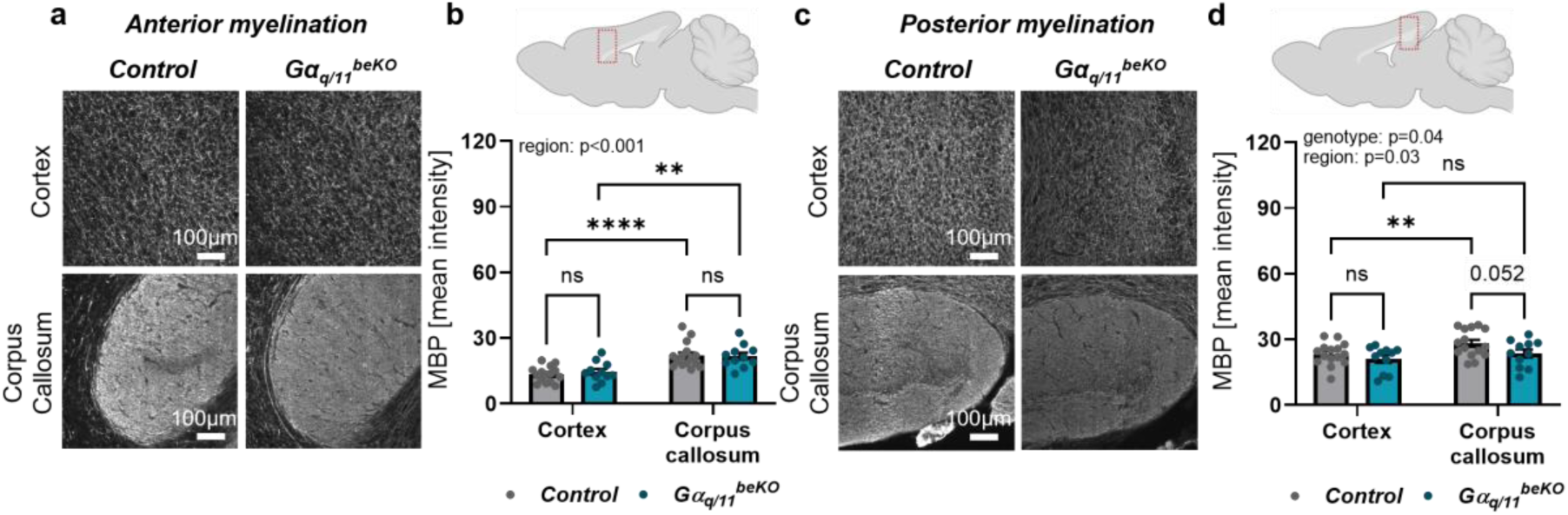
***Aged Gα_q/11_^beKO^ mice structural alterations in the white matter of the brain.*** (**a**) Representative images of myelin basic protein (MBP) in anterior cortex and corpus callosum of control and Gα_q/11_^beKO^ mice are shown. (**b**) Upper illustration indicates the area of imaging and lower part shows the quantification of MBP intensities in respective areas of aged control and Gα_q/11_^beKO^ mice. N=10. Significantly different variable is indicated in the graph. (**c**) Representative images of MBP in posterior cortex and corpus callosum of control and Gα_q/11_^beKO^ mice are shown. (**d**) Upper illustration indicates the area of imaging and lower part shows the quantification of MBP intensities in respective areas of aged control and *Gα_q/11_^beKO^* mice. N=10. Significantly different variable is indicated in the graph. Shown are means±SEM. ns: p>0.05, **p<0.01, ****p<0.0001. Detailed information on the age and time after tamoxifen, as well as the exact test statistics and values, is provided in Supplemental Table 3.

### Vascular alterations in Gα_q/11_^beKO^ mice appear with age

To further investigate the role of the genes we deleted, we examined expression of *Gnaq* and *Gna11* in FACS-sorted brain endothelial cells from mice and found a stable expression at different ages (Supplemental Fig. 9a-c). This finding is supported by a single-cell study containing data from several cell types of young and aged mouse brains,^45^ which we re-analysed for *Gnaq* and *Gna11* (Supplemental Fig. 9d) using the single cell portal of the Broad Institute.^46^ Since the cognitive impairment in old Gα_q/11_^beKO^ mice has an endothelial origin, we investigated the vessel structure of Gα_q/11_^beKO^ mice at an older age. Indeed, we found an increased level of so-called string vessels – empty basement membrane tubes that indicate endothelial cell death^47^ – in all the areas we investigated (Fig. 6a, b). These string vessels were not related to a general upregulation of vascular basement membrane components such as collagen IV or laminin α5 (Supplemental Fig. 10a, b). The increase in string vessels was most apparent in the corpus callosum, and it was accompanied by a decrease in vessel density in this region but not in cortex or hippocampus (Fig. 6c, Supplemental Fig. 10c, d). This is notable given that the corpus callosum already has a lower vascular density than grey matter regions, and small vessels are discussed to contribute to white matter changes in ageing,^48^ thereby likely explaining the decreased myelination in this region in Gα_q/11_^beKO^ mice. Interestingly, the occurrence of string vessels was detectable in aged but not in young or midlife mice (Fig. 6d). Since ageing is associated with pericytic changes,^49^ oxidative stress,^5^ and the occurrence of senescence in endothelial cells,^50^ we looked for those alterations in aged Gα_q/11_^beKO^ mice. We found a tendency to reduced pericyte coverage in the cortex but not in the corpus callosum (Supplemental Fig. 10e), suggesting different processes leading to endothelial cell loss or pericyte alterations depending on the brain region. Vascular senescence and oxidative stress are strongly related,^50,51^ but whether a loss of endothelial Gα_q/11_ signalling affects these processes is unknown. To examine the general state of senescence in the brain tissue, we performed expression analysis of p21, a key senescence marker. We found that aged Gα_q/11_^beKO^ mice have a higher expression of p21 compared to controls (Fig. 6e), which was not the case in the brains of 1-year-old mice (Supplemental Fig. 10f). To further confirm that finding, we stained brain sections for phosphorylated γH2AX (pγH2AX), a marker that is increased in cell nuclei with dsDNA breaks and another indicator of senescent cells.^52^ Indeed, we found increased pγH2AX intensity in endothelial cell nuclei of aged Gα_q/11_^beKO^ mice (Fig. 6f, g) but no genotype effect in parenchymal nuclei (Supplemental Fig. 10g), suggesting an accelerated senescence program, particularly in cells lacking Gα_q/11_ signalling. Related to that, we found increased intensity of the oxidative stress marker 4-HNE in vessels of aged Gα ^beKO^ compared to control mice (Fig. 6h, i). Both markers, pγH2AX and 4-HNE, were not different in younger Gα ^beKO^ compared to control mice (Supplemental Fig. 10h, i).

**Figure 6:**
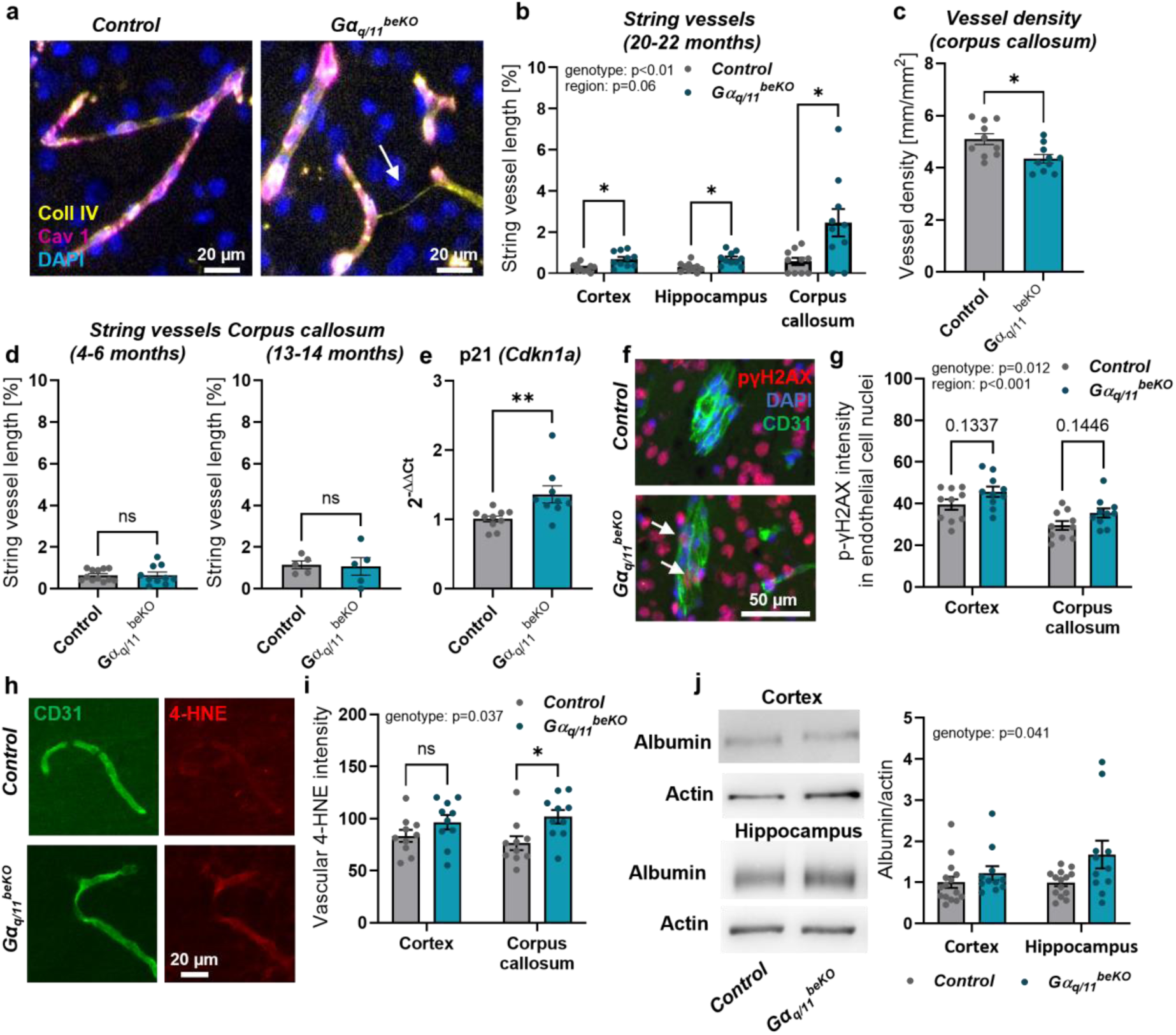
***Aged Gα_q/11_^beKO^ mice show vascular alterations in the white matter of the brain.*** (**a**) Representative images of staining for endothelial cells (caveolin 1, Cav 1) and basement membranes (collagen IV, Coll IV) in the cortex of control and Gα_q/11_^beKO^ mice are shown. Arrow indicates a string vessel, an empty basement membrane tube, indicating loss of endothelial cells. (**b**) Quantification of string vessels in different regions of the brains of aged control and Gα_q/11_^beKO^ mice. N=9-10. Significantly different variables are indicated in the graph. (**c**) Quantification of vessel density in the corpus callosum of aged control and Gα_q/11_^beKO^ mice. N=9-10. (**d**) Quantification of string vessels in the corpus callosum control and Gα_q/11_^beKO^ mice at different ages. N=5-12. (**e**) Quantification of the mRNA expression of the senescence marker p21 in brain tissue from aged control and Gα_q/11_^beKO^ mice. (**f**, **g**) Representative images with arrows indicating phosphorylated γH2AX in endothelial cells (f) and quantification thereof (g) of the cortex and corpus callosum from aged control and Gα_q/11_^beKO^ mice. N=10. Significantly different variables are indicated in the graph. (**h**, **i**) Representative images (h) and quantification (i) of the oxidative stress marker 4-HNE in the vessels of the cortex and corpus callosum from aged control and Gα_q/11_^beKO^ mice. N=10. Significantly different variables are indicated in the graph. (**j**) Representative images and quantification of Western blots of albumin in cortical and hippocampal brain tissue of aged control and *Gα_q/11_^beKO^* mice. N=11-15. Significantly different variable is indicated in the graph. Shown are means±SEM. ns: p>0.05, *p<0.05, **p<0.01, ***p<0.001. Detailed information on the age and time after tamoxifen, as well as the exact test statistics and values, is provided in Supplemental Table 3.

In addition to the vascular deficit, we also found a slight increase in BBB permeability in the hippocampus but not the cortex, measured by albumin content in brain tissue of aged perfused Gα ^beKO^ mice (Fig. 6j). Again, this vascular pathology was not present in younger mice (Supplemental Fig. 10j, k). We did not find a difference in occludin, ZO-1, or claudin 5 expression (Supplemental Fig. 10l-n), making other mechanisms than tight junction interruptions likely to be the cause for an increased permeability.

### Gα_q/11_ in brain endothelial cells interferes with VEGF signalling

Gα_q/11_ signalling in endothelial cells is related not only to NO release but also to signalling induced by VEGF, a vasoprotective agent.^53^ The loss of endothelial cells in the brain might be caused by reduced VEGF signalling, although we did not observe any differences in plasma VEGF levels between control and Gα_q/11_^beKO^ mice (Supplemental Fig. 11a). However, the protein levels of the main VEGF receptor interacting with Gα_q/11_ signalling, VEGFR2, were reduced in the brains of aged Gα_q/11_^beKO^ mice (Fig. 7a), independently of mRNA expression (Supplemental Fig. 11b). Since we were not able to measure the phosphorylated version of the VEGFR2 or downstream signalling molecules in brain tissue, we performed *in vitro* experiments in primary brain endothelial cells. By doing so, we found the VEGF-A-induced intracellular transduction being altered by impaired Gα_q/11_ signalling. While autophosphorylation of the VEGFR2 and phosphorylation of ERK1/2 were not affected in primary brain endothelial cells of Gα_q/11_^beKO^ mice (Fig. 7e-g), the VEGF-induced phosphorylation of eNOS (Fig. 7e, h) and the intracellular calcium increase were diminished upon loss of Gα_q/11_ signalling (Fig. 7b-d), indicating an interaction downstream of the VEGFR2 affecting NO production, although the baseline expression of the VEGFR2 was not altered in isolated primary brain endothelial cells (Fig. 7i). Since many regressing blood vessels in the brain were recently shown to regrow again,^54^ an impaired VEGFR2-related vessel proliferation might contribute to the increased occurrence of string vessels in those mice. Disturbed vascularisation and impaired VEGF signalling in the whole body are strongly correlated with a decreased lifespan and regression of organ vasculature.^55^ If the brain and its vasculature plays an essential role in determining life expectancy, this effect could help explain the increased mortality rate observed in Gα ^beKO^ mice in the course of ageing (Supplemental Fig. 11b), especially since peripheral organs and body weight were unaffected in Gα ^beKO^ mice during ageing (Supplemental Fig. 11c, d).

**Figure 7:**
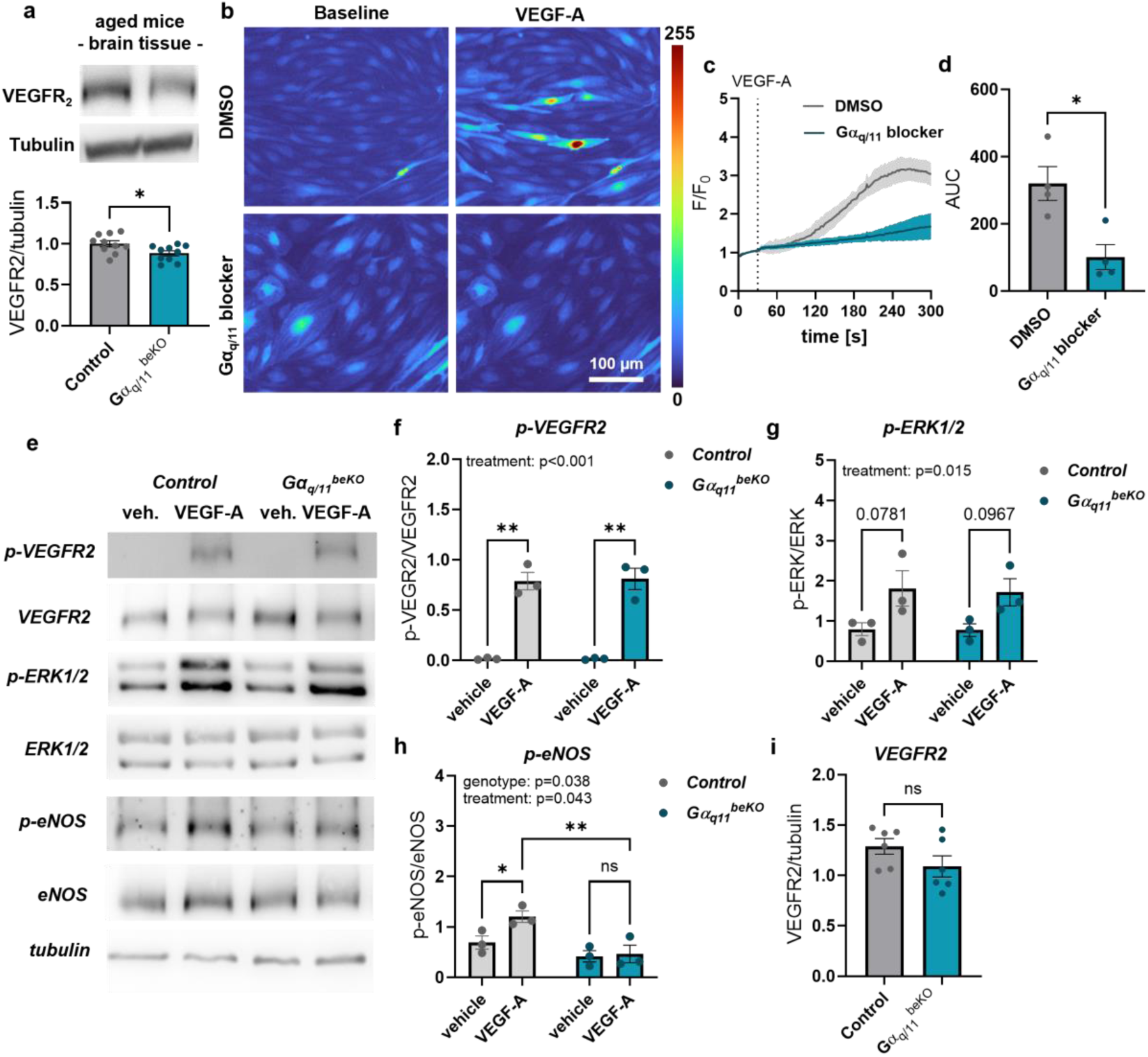
***Gα_q/11_ in brain endothelial cells interferes with VEGF signalling.*** (**a**) Representative images and quantification of Western blots of VEGFR2 in cortical brain tissue of aged control and Gα_q/11_^beKO^ mice. Student’s *t*-test. N=10. (**b-d**) Representative images (b) and quantification (c, d) of the calcium response in primary brain endothelial cells from *Cdh5*-GCaMP8 mice to VEGF (50 ng/mL) exposure with DMSO (vehicle) or the Gα_q/11_ blocker FR900359 (10 µM). N=4. (**e-i**) Representative images (d) and quantification (f-h) of VEGFR2 (i) and phosphorylation of VEGFR2 (f), ERK1/2 (g), and eNOS (h) in primary brain endothelial cells prepared from Gα_q/11_^beKO^ mice after tamoxifen treatment. Cells were stimulated with VEGF (50 ng/mL) or vehicle (medium) for 10 min before lysis. N=3. Significantly different variables are indicated in the graphs. Shown are means±SEM. ns: p>0.05, *p<0.05, **p<0.01, ****p<0.0001. Detailed information on the age and time after tamoxifen, as well as the exact test statistics and values, is provided in Supplemental Table 3.

## DISCUSSION

In summary, we present a mouse model of an inducible and isolated cerebral endothelial dysfunction that leads to an impaired cerebrovascular reactivity to CO_2_ and neuronal activity in adult mice. As they age, these mice develop mild cognitive impairment, indicating a causal relationship between endothelial dysfunction and cognitive deficits. While many diseases lead to endothelial dysfunction and cognitive deficits, it is unclear whether endothelial dysfunction itself contributes to impaired brain function. Here, we show that reduced endothelial GPCR signalling and vascular reactivity result in increased tau phosphorylation in the cortex and the hippocampus. This effect could be explained by reduced spontaneous and stimulated NO release from endothelial cells. Endothelium-derived mediators such as NO and VEGF affect not only the vasculature but are also essential for neuronal function.^56^ Reduced expression of VEGFR2 in the brains of these mice and altered VEGF-induced intracellular signalling in brain endothelial cells suggest a local imbalance of factors affecting vascular as well as neuron-mediated functions. Based on neuronal and synaptic markers, we demonstrate that neurons of Gα_q/11_^beKO^ mice are not morphologically but functionally impaired, leading to reduced memory retrieval in aged mice. The combination of old age and impaired cerebrovascular reactivity leads to an increased occurrence of string vessels in various regions and capillary rarefaction in the white matter of the brain. String vessels, endothelial senescence, and vascular oxidative stress due to the loss of endothelial Gα_q/11_ signalling were most prevalent in the white matter of the brain in aged mice, suggesting that these areas are particularly susceptible to vascular dysfunction. The reduced myelin content may partially explain the memory impairment in these mice, since a similar phenomenon is discussed to contribute to cognitive impairment in patients with small vessel disease,^57^ and loss of white matter during ageing contributes to cognitive decline.^58^ Interestingly, the white matter is severely affected in cerebral small vessel diseases,^2,59^ and string vessels are a feature of other models of cerebral small vessel diseases as well.^25,60,61^ Recently, the capillaries at the interface between the grey and white matter of the brain were discussed to be of utmost importance for the blood drainage of the white matter, and hypoperfusion in these areas could lead to demyelination.^62^ Our findings of the capillary rarefaction in the white matter and the reduced diameter of venules in Gα_q/11_^beKO^ mice could hint at a deficit in blood drainage, especially in these susceptible regions important for neuronal communication.

Increased mortality in Gα_q/11_^beKO^ mice during ageing may be explained by vascular rarefaction and increased blood-brain barrier permeability, which could lead to epileptic seizures or ischemic events as seen in cSVD.^4^ We showed that VEGF-induced signalling in brain endothelial cells is partially dependent on Gα_q/11_, and VEGFR2 expression was reduced in aged Gα_q/11_^beKO^ mice, which may be critical for the development of string vessels, as it has been shown to affect the survival of physiologically obstructed capillaries and thereby prevent pruning.^63^ Since VEGF and Gα_q/11_ signalling are closely connected, either via direct interaction^53^ or by phospholipase C interactions,^64^ the loss of the VEGF-induced calcium increase and eNOS activation without functional Gα_q/11_ signalling decreases NO availability even more, thereby contributing to endothelial dysfunction. The reduced NO release at baseline in Gα_q/11_-inhibited endothelial cells might indicate a baseline activity of Gα_q/11_ proteins, leading to constant eNOS activation and NO release. Since a reduced NO availability and a shift to reactive oxygen species are hallmarks of endothelial dysfunction, the Gα_q/11_ proteins may be a key part of maintaining the NO level in a physiological range. Future studies with either supplementing VEGF receptor agonists or NO donors would clarify the causal relation between memory deficits and cerebral endothelial dysfunction due to loss of Gα_q/11_ in the brain endothelium.

The limitations of this study include the use of a transgenic model system of cerebral endothelial dysfunction with limited translational significance. However, together with future studies mentioned above, we provide a potential targetable mechanism towards a translational implementation in the clinical treatment of patients suffering from cSVD, although a local treatment of the brain vasculature without affecting the periphery is still a challenge. In addition, single-cell data of brain tissue from cSVD patients would be interesting to examine the VEGF- and Gα_q/11_-related pathways in different parts of the neurovasculature, since we were describing mouse brain endothelial cells only.

Gα_q/11_^beKO^ mice provide a new model for investigating age-induced and endothelium-mediated cognitive impairment, making them ideal for exploring the mechanisms and targets of neurovascular dysfunction prevention in the elderly. It reflects findings known from cSVD and might be suitable to examine endothelium-caused effects in the non-amyloid form of this disease. We identified the endothelial Gα_q/11_ signalling pathway as a protective mechanism for cerebrovascular reactivity and the ageing vasculature in the brain. Microvascular targeting in endothelial dysfunction could counteract age-related neurological disorders, as demonstrated by blood-brain barrier-stabilising approaches^65,66^ or by VEGF administration.^55^ In this study, we present endothelial Gα_q/11_ signalling as an additional protective pathway that should be maintained and could be targeted in cerebrovascular dysfunction.

## CONTRIBUTORS

D.S., L.K., P.E., D.Z., R.N., N.N., C.E.H., A-S.G., A.M., J.S., P.L., B.L., and J.W. conducted experiments and analysed experimental data. G.A.C. performed angiography analyses. J.F., G.H., T.A.L., Ü.Ö., E.K., S.O., N.W., E.F., and M.S. provided essential resources. R.No., V.P., E.F., H.M.-F., S.O., and N.W. helped with data analysis and discussion. D.S., M.S., and J.W. designed the research. J.W. supervised the project and wrote the manuscript with editing and review by all authors.

## ACKNOWLEDGMENTS

We would like to thank Ines Stölting (University of Lübeck, Germany) for her expert technical assistance. We also thank Sonja Binder (University of Lübeck, Germany) for support with behavioural tests and statistical support. We thank Michael I. Kotlikoff (Cornell University, Ithaca, US) for providing *Cdh5*-GCaMP8 mice. Graphical inlets were created with BioRender.com.

## DATA SHARING STATEMENT

The data that support the findings of this study are available from the corresponding author upon reasonable request.

## REFERENCES

1. Li T, Huang Y, Cai W, et al. Age-related cerebral small vessel disease and inflammaging. Cell Death Dis. Oct 30 2020;11(10):932. doi:10.1038/s41419-020-03137-x

2. Markus HS, Joutel A. The pathogenesis of cerebral small vessel diseases and vascular cognitive impairment. Physiological reviews. Feb 18 2025;doi:10.1152/physrev.00028.2024

3. Debette S, Schilling S, Duperron MG, Larsson SC, Markus HS. Clinical Significance of Magnetic Resonance Imaging Markers of Vascular Brain Injury: A Systematic Review and Meta-analysis. JAMA Neurol. Jan 1 2019;76(1):81–94. doi:10.1001/jamaneurol.2018.3122

4. Kremer R, Williams A, Wardlaw J. Endothelial cells as key players in cerebral small vessel disease. Nature reviews Neuroscience. Mar 2025;26(3):179–188. doi:10.1038/s41583-024-00892-0

5. Chen MB, Yang AC, Yousef H, et al. Brain Endothelial Cells Are Exquisite Sensors of Age-Related Circulatory Cues. Cell reports. Mar 31 2020;30(13):4418–4432 e4. doi:10.1016/j.celrep.2020.03.012

6. Bennett HC, Zhang Q, Wu YT, et al. Aging drives cerebrovascular network remodeling and functional changes in the mouse brain. Nature communications. Jul 30 2024;15(1):6398. doi:10.1038/s41467-024-50559-8

7. Siragusa M, Fleming I. The eNOS signalosome and its link to endothelial dysfunction. Pflugers Archiv : European journal of physiology. Jul 2016;468(7):1125–1137. doi:10.1007/s00424-016-1839-0

8. Liao FF, Lin G, Chen X, et al. Endothelial Nitric Oxide Synthase-Deficient Mice: A Model of Spontaneous Cerebral Small-Vessel Disease. The American journal of pathology. Nov 2021;191(11):1932–1945. doi:10.1016/j.ajpath.2021.02.022

9. Huang PL, Huang Z, Mashimo H, et al. Hypertension in mice lacking the gene for endothelial nitric oxide synthase. Nature. Sep 21 1995;377(6546):239–42. doi:10.1038/377239a0

10. Harraz OF, Longden TA, Hill-Eubanks D, Nelson MT. PIP2 depletion promotes TRPV4 channel activity in mouse brain capillary endothelial cells. eLife. Aug 7 2018;7doi:10.7554/eLife.38689

11. Harraz OF, Longden TA, Dabertrand F, Hill-Eubanks D, Nelson MT. Endothelial GqPCR activity controls capillary electrical signaling and brain blood flow through PIP2 depletion. Proceedings of the National Academy of Sciences of the United States of America. Apr 10 2018;115(15):E3569–E3577. doi:10.1073/pnas.1800201115

12. Hashad AM, Abd-Alhaseeb MM, Lim XR, Mathieu NM, Harraz OF. PIP(2) corrects an endothelial Piezo1 channelopathy. Proceedings of the National Academy of Sciences of the United States of America. Dec 30 2025;122(52):e2522750122. doi:10.1073/pnas.2522750122

13. Dabertrand F, Harraz OF, Koide M, et al. PIP(2) corrects cerebral blood flow deficits in small vessel disease by rescuing capillary Kir2.1 activity. Proceedings of the National Academy of Sciences of the United States of America. Apr 27 2021;118(17)doi:10.1073/pnas.2025998118

14. Wang S, Iring A, Strilic B, et al. P2Y(2) and Gq/G(1)(1) control blood pressure by mediating endothelial mechanotransduction. Research Support, Non-U.S. Gov’t. The Journal of clinical investigation. Aug 3 2015;125(8):3077–86. doi:10.1172/JCI81067

15. Wenzel J, Hansen CE, Bettoni C, et al. Impaired endothelium-mediated cerebrovascular reactivity promotes anxiety and respiration disorders in mice. Proceedings of the National Academy of Sciences of the United States of America. Jan 21 2020;117(3):1753–1761. doi:10.1073/pnas.1907467117

16. Ridder DA, Lang MF, Salinin S, et al. TAK1 in brain endothelial cells mediates fever and lethargy. Research Support, Non-U.S. Gov’t. The Journal of experimental medicine. Dec 19 2011;208(13):2615–23. doi:10.1084/jem.20110398

17. Wettschureck N, Rutten H, Zywietz A, et al. Absence of pressure overload induced myocardial hypertrophy after conditional inactivation of Galphaq/Galpha11 in cardiomyocytes. Research Support, Non-U.S. Gov’t. Nature medicine. Nov 2001;7(11):1236–40. doi:10.1038/nm1101-1236

18. Mori T, Tanaka K, Buffo A, Wurst W, Kuhn R, Gotz M. Inducible gene deletion in astroglia and radial glia--a valuable tool for functional and lineage analysis. Glia. Jul 2006;54(1):21–34. doi:10.1002/glia.20350

19. Madisen L, Zwingman TA, Sunkin SM, et al. A robust and high-throughput Cre reporting and characterization system for the whole mouse brain. Nature neuroscience. Jan 2010;13(1):133–40. doi:10.1038/nn.2467

20. Körbelin J DG, Michelfelder S, Ridder D A, Hunger A, Wenzel J, Seismann H, Lampe M, Bannach J, Pasparakis M, Kleinschmidt J A, Schwaninger M, Trepel M. A brain microvasculature endothelial cell-specific viral vector with the potential to treat neurovascular and neurological diseases. EMBO Molecular Medicine 2016;8(6):609–25.

21. Lee FK, Lee JC, Shui B, et al. Genetically engineered mice for combinatorial cardiovascular optobiology. eLife. Oct 29 2021;10doi:10.7554/eLife.67858

22. Longden TA, Mughal A, Hennig GW, et al. Local IP(3) receptor-mediated Ca(2+) signals compound to direct blood flow in brain capillaries. Science advances. Jul 2021;7(30)doi:10.1126/sciadv.abh0101

23. Huber G, Ogrodnik M, Wenzel J, et al. Telmisartan prevents high-fat diet-induced neurovascular impairments and reduces anxiety-like behavior. Journal of cerebral blood flow and metabolism : official journal of the International Society of Cerebral Blood Flow and Metabolism. Sep 2021;41(9):2356–2369. doi:10.1177/0271678X211003497

24. Csaszar E, Lenart N, Cserep C, et al. Microglia modulate blood flow, neurovascular coupling, and hypoperfusion via purinergic actions. The Journal of experimental medicine. Mar 7 2022;219(3)doi:10.1084/jem.20211071

25. Ridder DA, Wenzel J, Muller K, et al. Brain endothelial TAK1 and NEMO safeguard the neurovascular unit. The Journal of experimental medicine. Sep 21 2015;212(10):1529–49. doi:10.1084/jem.20150165

26. Binder S, Baier PC, Molle M, Inostroza M, Born J, Marshall L. Sleep enhances memory consolidation in the hippocampus-dependent object-place recognition task in rats. Neurobiol Learn Mem. Feb 2012;97(2):213–9. doi:10.1016/j.nlm.2011.12.004

27. Murai T, Okuda S, Tanaka T, Ohta H. Characteristics of object location memory in mice: Behavioral and pharmacological studies. Physiol Behav. Jan 30 2007;90(1):116–24. doi:10.1016/j.physbeh.2006.09.013

28. Binder S, Molle M, Lippert M, et al. Monosynaptic Hippocampal-Prefrontal Projections Contribute to Spatial Memory Consolidation in Mice. The Journal of neuroscience : the official journal of the Society for Neuroscience. 2019;39(35):6978–6991.

29. Wenzel J, Lampe J, Muller-Fielitz H, et al. The SARS-CoV-2 main protease M(pro) causes microvascular brain pathology by cleaving NEMO in brain endothelial cells. Nature neuroscience. Nov 2021;24(11):1522–1533. doi:10.1038/s41593-021-00926-1

30. Assmann JC, Muller K, Wenzel J, et al. Isolation and Cultivation of Primary Brain Endothelial Cells from Adult Mice. Bio-protocol. May 20 2017;7(10)doi:10.21769/BioProtoc.2294

31. Schrage R, Schmitz AL, Gaffal E, et al. The experimental power of FR900359 to study Gq-regulated biological processes. Nature communications. 2015;6:10156. doi:10.1038/ncomms10156

32. Spyropoulos D, Wenzel J. Live Calcium Imaging on Mouse Primary Brain Endothelial Cells. Methods Mol Biol. 2025;2956:119–124. doi:10.1007/978-1-0716-4706-6_11

33. Nizar K, Uhlirova H, Tian P, et al. In vivo stimulus-induced vasodilation occurs without IP3 receptor activation and may precede astrocytic calcium increase. Research Support, N.I.H., Extramural Research Support, Non-U.S. Gov’t. The Journal of neuroscience : the official journal of the Society for Neuroscience. May 8 2013;33(19):8411–22. doi:10.1523/JNEUROSCI.3285-12.2013

34. Institoris A, Vandal M, Peringod G, et al. Astrocytes amplify neurovascular coupling to sustained activation of neocortex in awake mice. Nature communications. Dec 22 2022;13(1):7872. doi:10.1038/s41467-022-35383-2

35. Mughal A, Hennig GW, Heppner T, Tsoukias NM, Hill-Eubanks D, Nelson MT. Electrocalcium coupling in brain capillaries: Rapidly traveling electrical signals ignite local calcium signals. Proceedings of the National Academy of Sciences of the United States of America. Dec 17 2024;121(51):e2415047121. doi:10.1073/pnas.2415047121

36. Schaeffer S, Iadecola C. Revisiting the neurovascular unit. Nature neuroscience. Sep 2021;24(9):1198–1209. doi:10.1038/s41593-021-00904-7

37. de Leeuw FE, de Kleine M, Frijns CJ, Fijnheer R, van Gijn J, Kappelle LJ. Endothelial cell activation is associated with cerebral white matter lesions in patients with cerebrovascular disease. Annals of the New York Academy of Sciences. Nov 2002;977:306–14. doi:10.1111/j.1749-6632.2002.tb04831.x

38. Rajani RM, Quick S, Ruigrok SR, et al. Reversal of endothelial dysfunction reduces white matter vulnerability in cerebral small vessel disease in rats. Sci Transl Med. Jul 4 2018;10(448)doi:10.1126/scitranslmed.aam9507

39. Vlegels N, van den Brink H, Kopczak A, et al. The relation between cerebral small vessel function and white matter microstructure in monogenic and sporadic small vessel disease - the ZOOM@SVDs study. Cereb Circ Cogn Behav. 2025;8:100383. doi:10.1016/j.cccb.2025.100383

40. Faraco G, Hochrainer K, Segarra SG, et al. Dietary salt promotes cognitive impairment through tau phosphorylation. Nature. Oct 2019;574(7780):686–690. doi:10.1038/s41586-019-1688-z

41. Wang Y, Mandelkow E. Tau in physiology and pathology. Nature reviews Neuroscience. Jan 2016;17(1):5–21. doi:10.1038/nrn.2015.1

42. Bathina S, Das UN. Brain-derived neurotrophic factor and its clinical implications. Arch Med Sci. Dec 10 2015;11(6):1164–78. doi:10.5114/aoms.2015.56342

43. Quirie A, Mor D, Meloux A, et al. Anxio-depressive phenotype and impaired memory in mice with a conditional knockout of brain-derived neurotrophic factor in endothelial cells. Am J Physiol Cell Physiol. Jan 1 2025;328(1):C303–C314. doi:10.1152/ajpcell.00699.2024

44. Walsh J, Tozer DJ, Sari H, et al. Microglial activation and blood-brain barrier permeability in cerebral small vessel disease. Brain : a journal of neurology. Jun 22 2021;144(5):1361–1371. doi:10.1093/brain/awab003

45. Ximerakis M, Lipnick SL, Innes BT, et al. Single-cell transcriptomic profiling of the aging mouse brain. Nature neuroscience. Oct 2019;22(10):1696–1708. doi:10.1038/s41593-019-0491-3

46. Tarhan L, Bistline J, Chang J, Galloway B, Hanna E, Weitz E. Single Cell Portal: an interactive home for single-cell genomics data. bioRxiv. Jul 17 2023;doi:10.1101/2023.07.13.548886

47. Brown WR. A review of string vessels or collapsed, empty basement membrane tubes. Journal of Alzheimer’s disease : JAD. 2010;21(3):725–39. doi:10.3233/JAD-2010-100219

48. Salat DH. Imaging small vessel-associated white matter changes in aging. Neuroscience. Sep 12 2014;276:174–86. doi:10.1016/j.neuroscience.2013.11.041

49. Berthiaume AA, Schmid F, Stamenkovic S, et al. Pericyte remodeling is deficient in the aged brain and contributes to impaired capillary flow and structure. Nature communications. Oct 7 2022;13(1):5912. doi:10.1038/s41467-022-33464-w

50. Kiss T, Nyul-Toth A, Balasubramanian P, et al. Single-cell RNA sequencing identifies senescent cerebromicrovascular endothelial cells in the aged mouse brain. GeroScience. Apr 2020;42(2):429–444. doi:10.1007/s11357-020-00177-1

51. Real MGC, Falcione SR, Boghozian R, et al. Endothelial Cell Senescence Effect on the Blood-Brain Barrier in Stroke and Cognitive Impairment. Neurology. Dec 10 2024;103(11):e210063. doi:10.1212/WNL.0000000000210063

52. d’Adda di Fagagna F, Reaper PM, Clay-Farrace L, et al. A DNA damage checkpoint response in telomere-initiated senescence. Nature. Nov 13 2003;426(6963):194–8. doi:10.1038/nature02118

53. Sivaraj KK, Li R, Albarran-Juarez J, et al. Endothelial Galphaq/11 is required for VEGF-induced vascular permeability and angiogenesis. Cardiovascular research. Oct 1 2015;108(1):171–80. doi:10.1093/cvr/cvv216

54. Gao X, Chen XJ, Ye M, et al. Reduction of neuronal activity mediated by blood-vessel regression in the adult brain. Nature communications. Jul 1 2025;16(1):5840. doi:10.1038/s41467-025-60308-0

55. Grunewald M, Kumar S, Sharife H, et al. Counteracting age-related VEGF signaling insufficiency promotes healthy aging and extends life span. Science. Jul 30 2021;373(6554)doi:10.1126/science.abc8479

56. Aksan B, Mauceri D. Beyond vessels: unraveling the impact of VEGFs on neuronal functions and structure. J Biomed Sci. Mar 6 2025;32(1):33. doi:10.1186/s12929-025-01128-8

57. Denecke J, Dewenter A, Lee J, et al. Reduced myelin contributes to cognitive impairment in patients with monogenic small vessel disease. Alzheimer’s & dementia : the journal of the Alzheimer’s Association. May 2025;21(5):e70127. doi:10.1002/alz.70127

58. Gunning-Dixon FM, Brickman AM, Cheng JC, Alexopoulos GS. Aging of cerebral white matter: a review of MRI findings. Int J Geriatr Psychiatry. Feb 2009;24(2):109–17. doi:10.1002/gps.2087

59. Benisty S, Reyes S, Godin O, et al. White-matter lesions without lacunar infarcts in CADASIL. Journal of Alzheimer’s disease : JAD. 2012;29(4):903–11. doi:10.3233/JAD-2012-111784

60. Muller K, Courtois G, Ursini MV, Schwaninger M. New Insight Into the Pathogenesis of Cerebral Small-Vessel Diseases. Stroke. Feb 2017;48(2):520–527. doi:10.1161/STROKEAHA.116.012888

61. Tong XK, Hamel E. Simvastatin restored vascular reactivity, endothelial function and reduced string vessel pathology in a mouse model of cerebrovascular disease. Journal of cerebral blood flow and metabolism : official journal of the International Society of Cerebral Blood Flow and Metabolism. Mar 2015;35(3):512–20. doi:10.1038/jcbfm.2014.226

62. Stamenkovic S, Schmid F, Gurler G, et al. Impaired capillary-venous drainage contributes to gliosis and demyelination in mouse white matter during aging. Nature neuroscience. Aug 12 2025;doi:10.1038/s41593-025-02023-z

63. Reeson P, Choi K, Brown CE. VEGF signaling regulates the fate of obstructed capillaries in mouse cortex. eLife. Apr 26 2018;7doi:10.7554/eLife.33670

64. Bhattacharya R, Kwon J, Li X, et al. Distinct role of PLCbeta3 in VEGF-mediated directional migration and vascular sprouting. J Cell Sci. Apr 1 2009;122(Pt 7):1025–34. doi:10.1242/jcs.041913

65. Greene C, Hanley N, Reschke CR, et al. Microvascular stabilization via blood-brain barrier regulation prevents seizure activity. Nature communications. Apr 14 2022;13(1):2003. doi:10.1038/s41467-022-29657-y

66. Martin M, Vermeiren S, Bostaille N, et al. Engineered Wnt ligands enable blood-brain barrier repair in neurological disorders. Science. Feb 18 2022;375(6582):eabm4459. doi:10.1126/science.abm4459

