## Supplemental data for "Gα_q/11_ signalling counteracts endothelial dysfunction in the brain and protects cognition in aged mice"

**for**

Contains:

Supplemental Figures 1-11

Supplemental Methods

Supplemental Tables 1-3

Corresponding author:

Jan Wenzel

Institute of Experimental and Clinical Pharmacology and Toxicology, Center of Brain, Behavior and Metabolism (CBBM), University of Lübeck, Ratzeburger Allee 160, 23562 Lübeck, Germany (+49) 451-3101-7224

SUPPLEMENTAL FIGURES

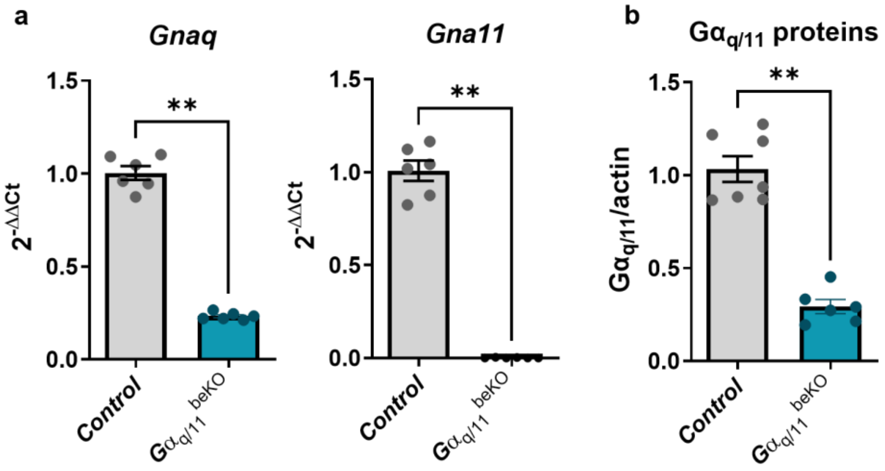

***Supplemental Figure 1: Successful deletion of G$\alpha_{q/11}$ in brain endothelial cells.*** (**a**) Gnaq and Gna11 mRNA expression in primary brain endothelial cells of G$\alpha_{q/11}^{beKO}$ and control mice. N=6. (**b**) Western blotting against G$\alpha_{q/11}$ proteins in primary brain endothelial cells prepared after tamoxifen treatment of G$\alpha_{q/11}^{beKO}$ and control mice. N=6-7. Shown are means±SEM.****p<0.0001. Detailed information on the age and time after tamoxifen, as well as the exact test statistics and values, is provided in Supplemental Table 3.

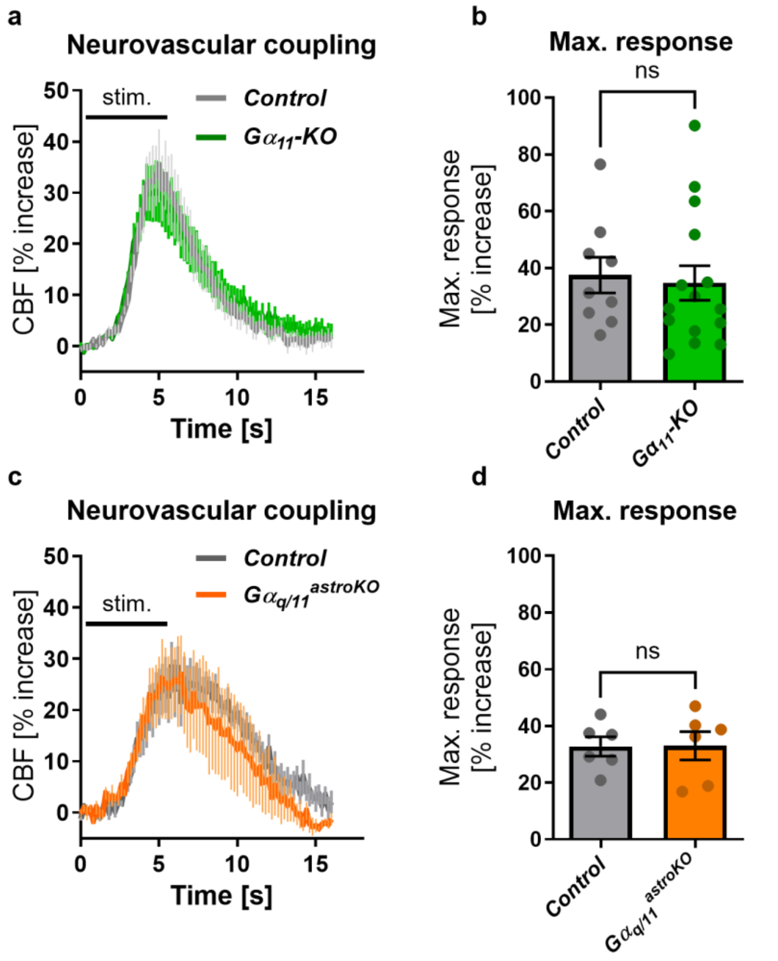

***Supplemental Figure 2: Whole genome deletion of Gna11 or astrocytic deletion of the $G\alpha_{q/11}$ signaling pathway does not affect neurovascular coupling.*** (**a**) Time course of cerebral blood flow (CBF) recording during whisker pad stimulation in $G\alpha_{11}$-KO and control mice. N=9-15. (**b**) Maximal CBF response of curves shown in (a). (**c**) Time course of CBF recording during whisker pad stimulation in $G\alpha_{q/11}^{astroKO}$ and control mice. N=6. (**d**) Maximal CBF response of curves shown in (c). Shown are means±SEM. ns: $p>0.05$, $p > 0.05$. Detailed information on the age and time after tamoxifen, as well as the exact test statistics and values, is provided in Supplemental Table 3.

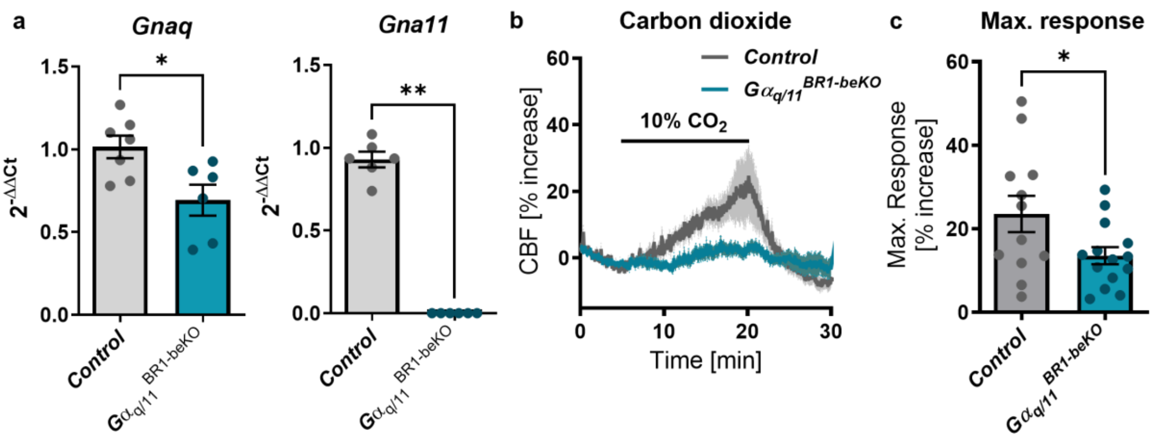

***Supplemental Figure 3: Brain endothelial deletion of Gα$_{q/11}$ by AAV-BR1-Cre expression impairs cerebrovascular reactivity.*** (**a**) *Gnaq* and *Gna11* mRNA expression in primary brain endothelial cells prepared after intravenous AAV injection ($1x10^{12}$ gp/mouse), either encoding a Cre recombinase (Gα$_{q/11}^{BR1\text{-}beKO}$) or a GFP (Control). N=6-7. (**b**) Time course of cerebral blood flow (CBF) recording during $CO_2$ stimulation. N=12-14. (**c**) Maximal CBF response of curves shown in (b). Shown are means±SEM. *p<0.05, **p<0.01. Detailed information on the age and time after tamoxifen, as well as the exact test statistics and values, is provided in Supplemental Table 3.

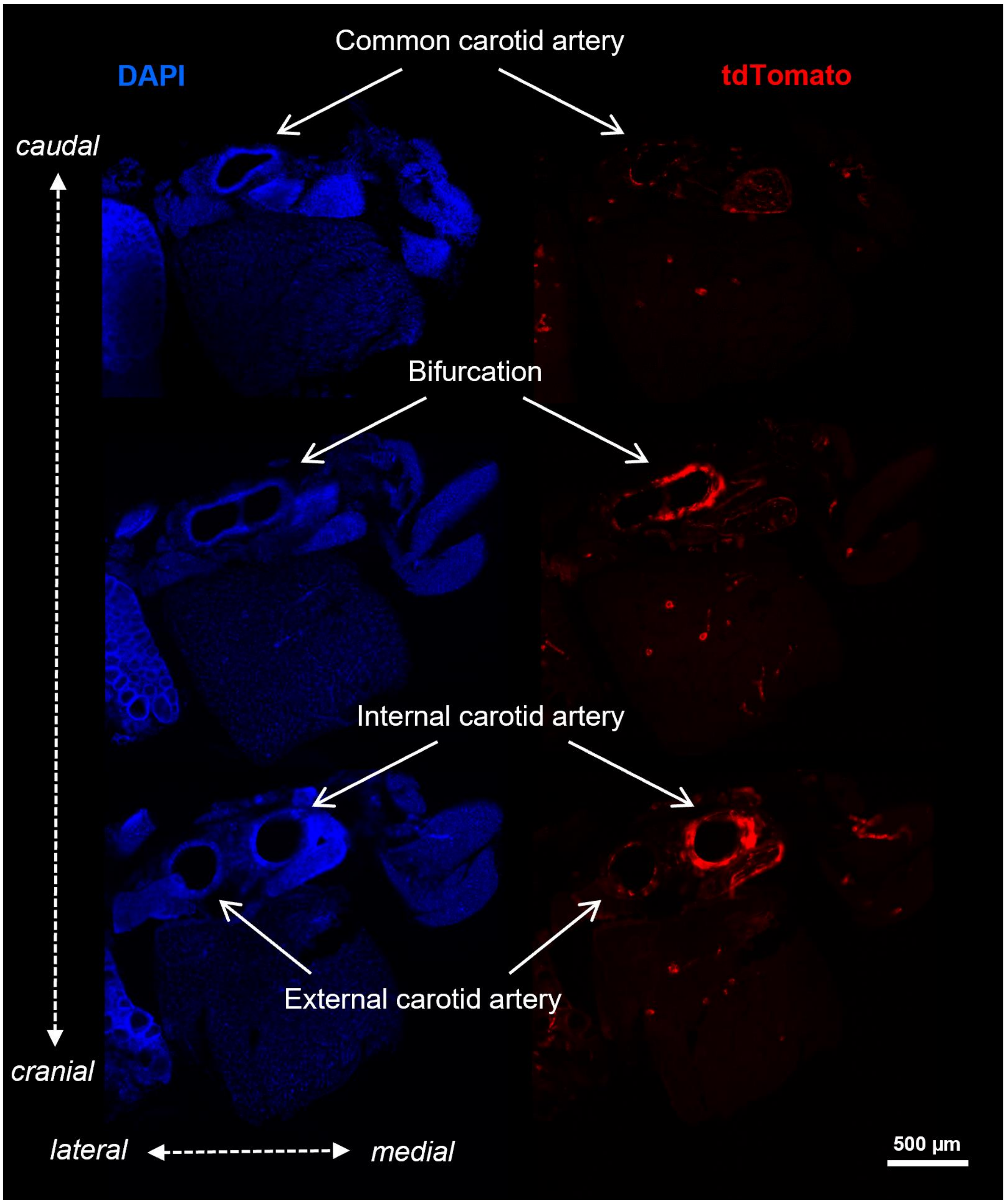

***Supplemental Figure 4: Recombination of the Slco1c1-CreER$^{T2}$ mouse line starts at the bifurcation between the internal and external carotid arteries.*** Shown are three different cross-sections of tissue in the region where the common carotid artery diverges into the internal and the external carotid arteries. A *Slco1c1*-CreER$^{T2}$; Ai14$^{+/fl}$ mouse, two weeks after the start of tamoxifen injection, was used. tdTomato indicates recombination, and DAPI was used to counterstain the tissue.

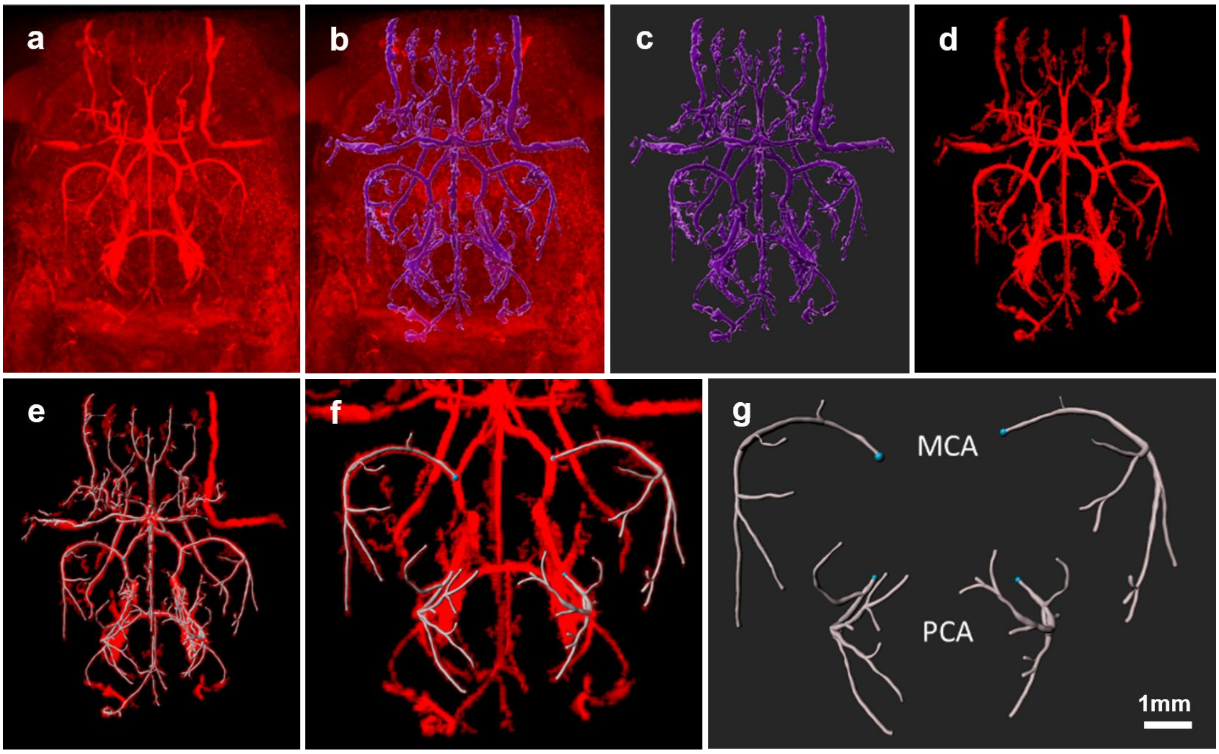

***Supplemental Figure 5: Workflow for processing TOF angiography data sets.*** An example of a data set obtained by time of flight (TOF) angiography MRI and its processing using the software Imaris is shown. (**a**) original data set, (**b**) surface creation overlay, (**c**) final surface creation after removing noise, (**d**) mask of the surface creation, (**e**) filaments detection, (**f**) magnification of (e) and selection of middle cerebral arteries (MCA) and posterior cerebral arteries (PCA) with the starting points (blue dots) at the circle of Willis. (**g**) shown are the final filaments used for quantification (see main Figure 2).

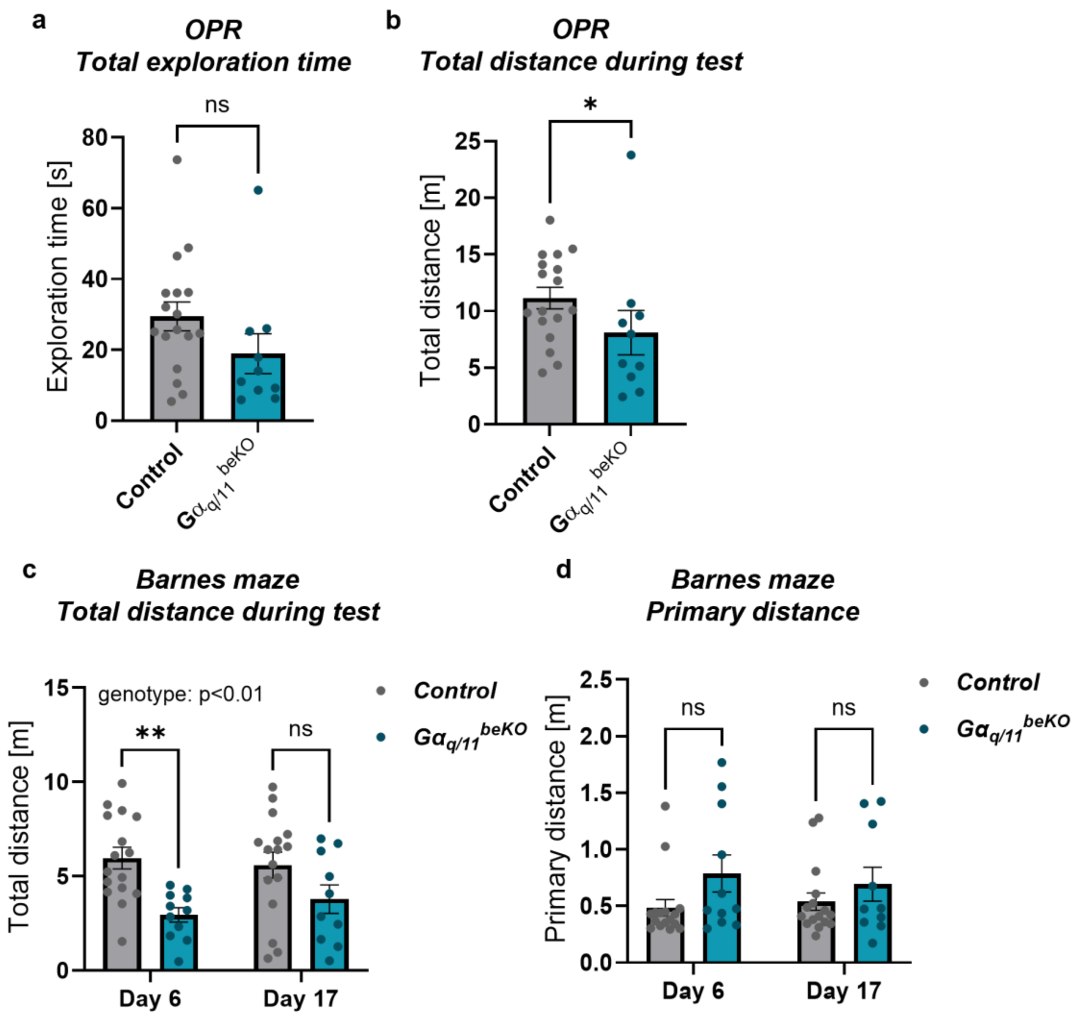

***Supplemental Figure 6: Motor activity during behaviour tests in aged $G\alpha_{q/11}^{beKO}$ and control mice.*** (**a**, **b**) Total exploration times and distances during the test of $G\alpha_{q/11}^{beKO}$ and control mice at the age of 18-20 months are shown. (**c, d**) Total and primary distances travelled by 20-22 months old $G\alpha_{q/11}^{beKO}$ and control mice during the test on days 6 and 17 are shown. N=11-16, Shown are means±SEM. ns: $p>0.05$, $^{**}p<0.01$. Detailed information on the age and time after tamoxifen, as well as the exact test statistics and values, is provided in Supplemental Table 3.

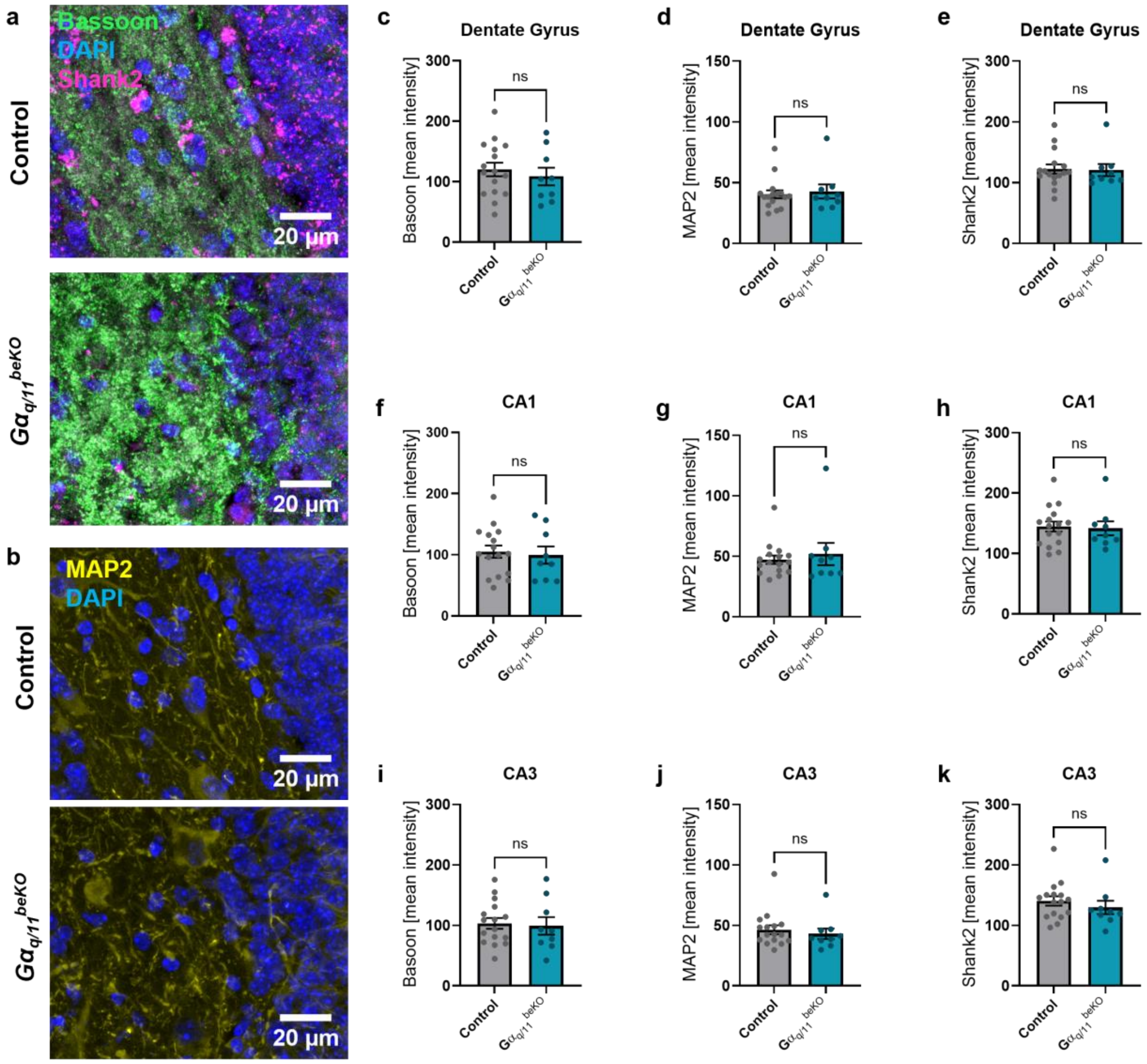

***Supplemental Figure 7: No difference in neuronal markers in hippocampal subregions of aged $G\alpha_{q/11}^{beKO}$ mice.*** (**a**, **b**) Representative images of staining for synaptic markers bassoon and shank2 (a) and structural marker MAP2 (b) in hippocampal tissue of 22-month-old control and $G\alpha_{q/11}^{beKO}$ mice. (**c-k**) Quantification of mean intensities of bassoon (c, f, i), MAP2 (d, g, j), and Shank2 (e, h, k) are shown for the hippocampal regions dentate gyrus (c-e), CA1 (f-h), and CA3 (i-k) in aged control and $G\alpha_{q/11}^{beKO}$ mice. Shown are means±SEM. Detailed information on the age and time after tamoxifen, as well as the exact test statistics and values, is provided in Supplemental Table 3.

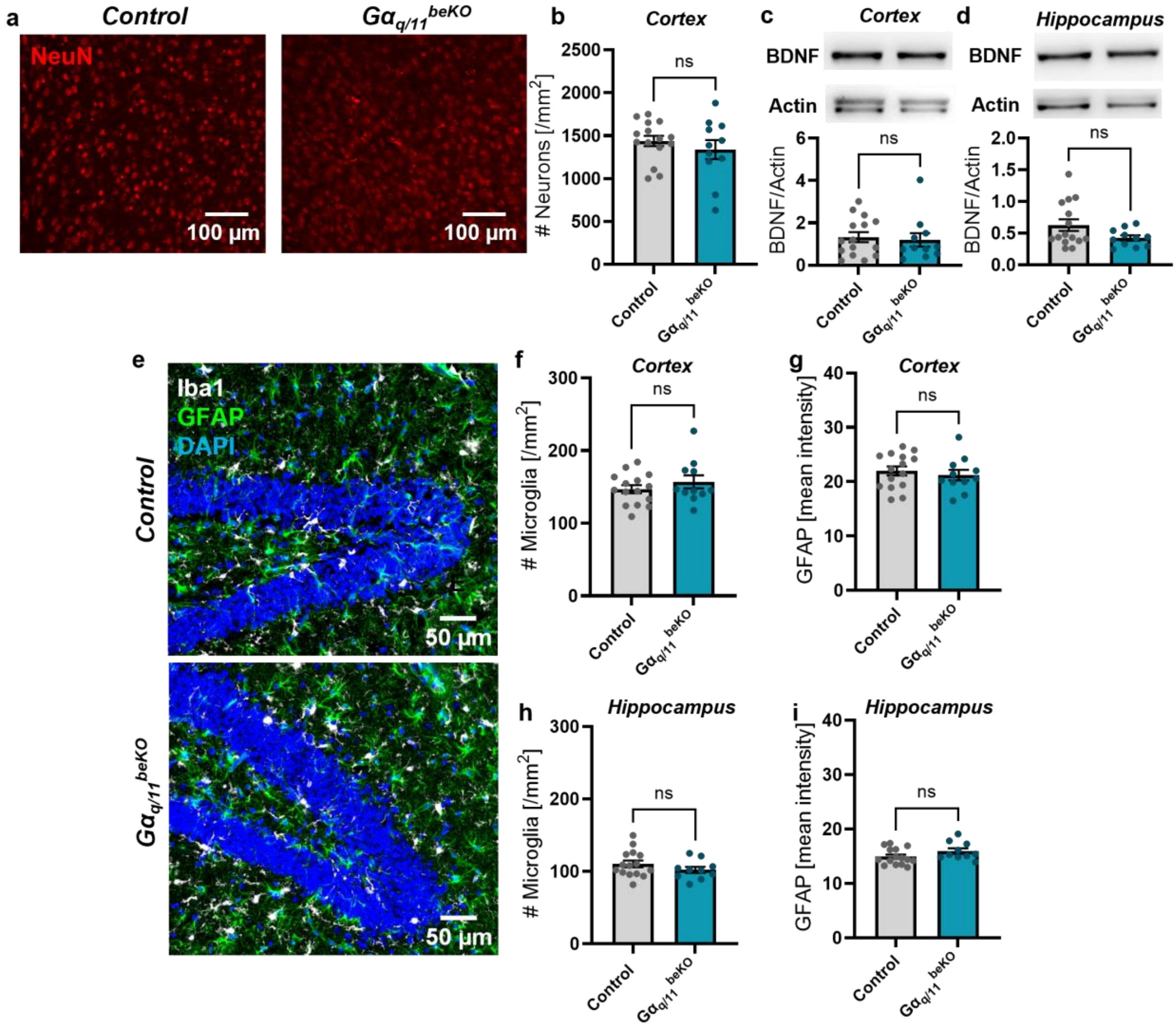

***Supplemental Figure 8: No difference in the number of neurons or neuroinflammatory markers in brains of aged $G\alpha_{q/11}^{beKO}$ mice.*** (**a**, **b**) Representative images (a) and quantification (b) of neuronal marker NeuN in cortex of 20-22-month-old control and $G\alpha_{q/11}^{beKO}$ mice. (**c**, **d**) Representative Western blots and quantification of brain-derived neurotrophic factor (BDNF) and actin in cortical (c) and hippocampal (d) tissue of aged control and $G\alpha_{q/11}^{beKO}$ mice are shown. (**e-i**) Representative images (e) and quantification (f-i) of microglia (Iba1; f, h) and astrocytic activation (GFAP, g, i) in cortical (f, g) and hippocampal (h, i) tissue of aged control and $G\alpha_{q/11}^{beKO}$ mice. N=10-15. Shown are means±SEM.ns: $p>0.05$. Detailed information on the age and time after tamoxifen, as well as the exact test statistics and values, is provided in Supplemental Table 3.

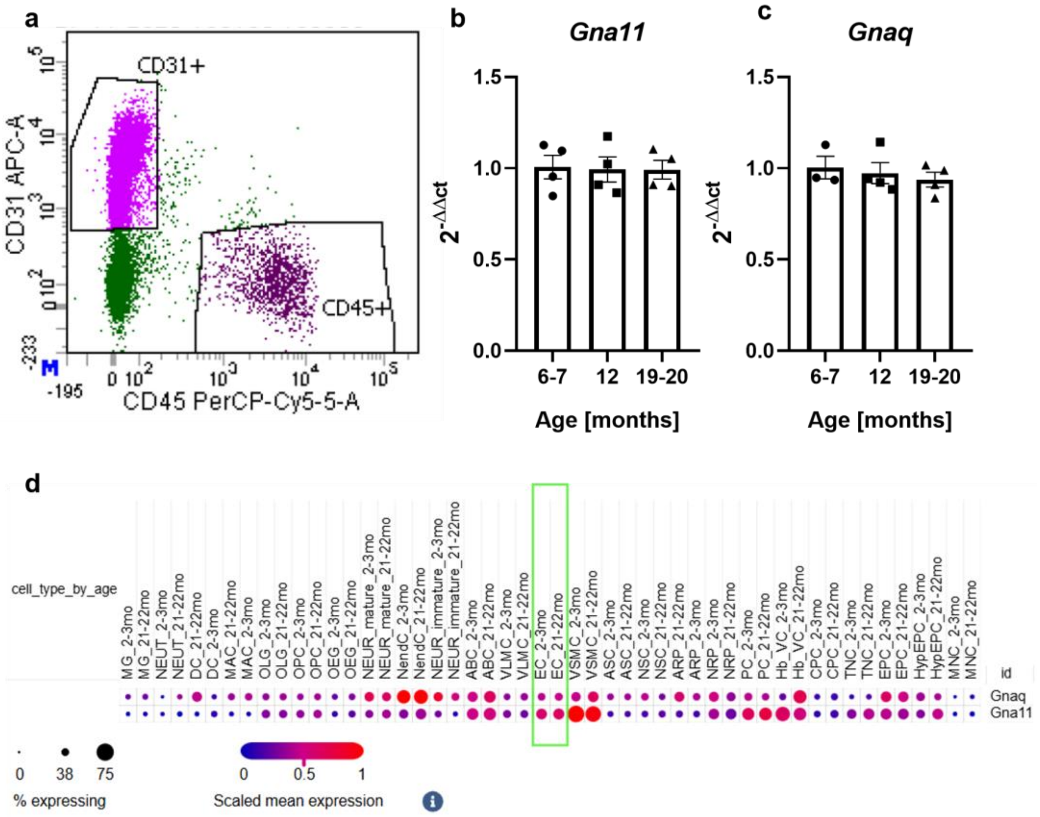

***Supplemental Figure 9: Gα$_{q/11}$ expression in aged mice.*** (**a-c**) Fluorescence-activated cell sorting (FACS) using CD31- and CD45-targeting antibodies was used to isolate endothelial cells ($CD31^{+}$, $CD45^{-}$) from the brains of mice at different ages. Shown are the gating strategy (a) and mRNA expression of *Gna11* (b) and *Gnaq* (c) in isolated cells. N=4. Shown are means±SEM. Detailed information on exact test statistics and values is provided in Supplemental Table 3. (**d**) Re-analysis of the expression of *Gnaq* and *Gna11* in a single-cell study on adult (2-3 months) and aged (21-22 months) brain cells published before (1) and accessed via the single-cell portal of the Broad Institute (2). The green bar indicates the endothelial cell cluster.

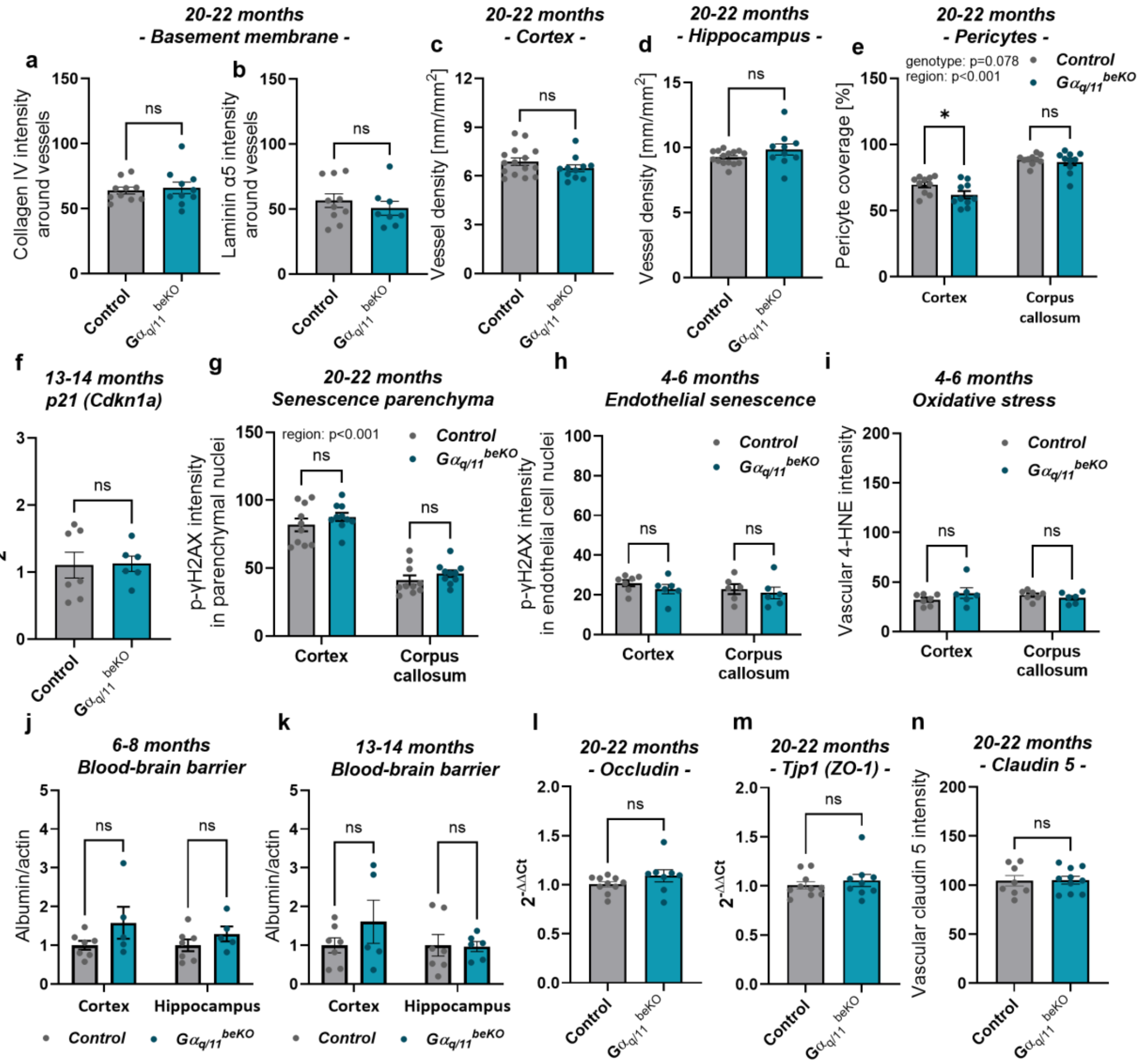

***Supplemental Figure 10: Vascular effects in aged $G\alpha_{q/11}^{beKO}$ mice.*** (**a, b**) Vascular extracellular matrix components collagen IV (a) and laminin α5 (b) were measured in the cerebral vasculature of 20-22-month-old control and $G\alpha_{q/11}^{beKO}$ mice by immunofluorescence staining. N=8-10. (**c, d**) Vessel densities in the cortical (c) and hippocampal (d) brain tissue of 20-22-month-old control and $G\alpha_{q/11}^{beKO}$ determined by immunofluorescence staining. N=8-10. (**e**) Pericytic coverage was determined by immunofluorescence staining for CD13 and endothelial cells in the cortex and corpus callosum of 20-22-month-old control and $G\alpha_{q/11}^{beKO}$ mice. N=8-10. Significantly different variables are indicated in the graph. (**f**) Expression of the senescence marker p21 in brain tissue of 13-14 months-old control and $G\alpha_{q/11}^{beKO}$ mice. N=6-7. (**g**) Senescence in the parenchyma of the cortex and the corpus callosum of 20-22-month-old control and $G\alpha_{q/11}^{beKO}$ mice, determined by staining for phosphorylated γH2AX and an endothelial cell marker. Only non-endothelial cell nuclei were analysed. Significantly different variable is indicated in the graph. N=8-10. (**h, i**) Senescence and oxidative stress in endothelial cells of the cortex and the corpus callosum of 4-6-month-old control and $G\alpha_{q/11}^{beKO}$ mice, determined by staining for phosphorylated γH2AX or 4-HNE and an endothelial cell marker. Only endothelial cells were analysed. N=5-7. (**j**, **k**) Examination of the blood-brain barrier permeability by Western blotting for albumin in cortical or hippocampal brain tissue from perfused 6-8-month-old (j) or 13-14-month-old (k) control and $G\alpha_{q/11}^{beKO}$ mice. N=5-7. (**l**, **m**, **n**) Expression of components of endothelial tight junctions by qPCR (l, occludin, and m, ZO-1) and immunofluorescent staining (n, claudin 5) in brain tissue of 20-22-month-old $G\alpha_{q/11}^{beKO}$ control mice. N=9-10. Shown are means±SEM. ns: p>0.05. Detailed information on the age and time after tamoxifen, as well as the exact test statistics and values, is provided in Supplemental Table 3.

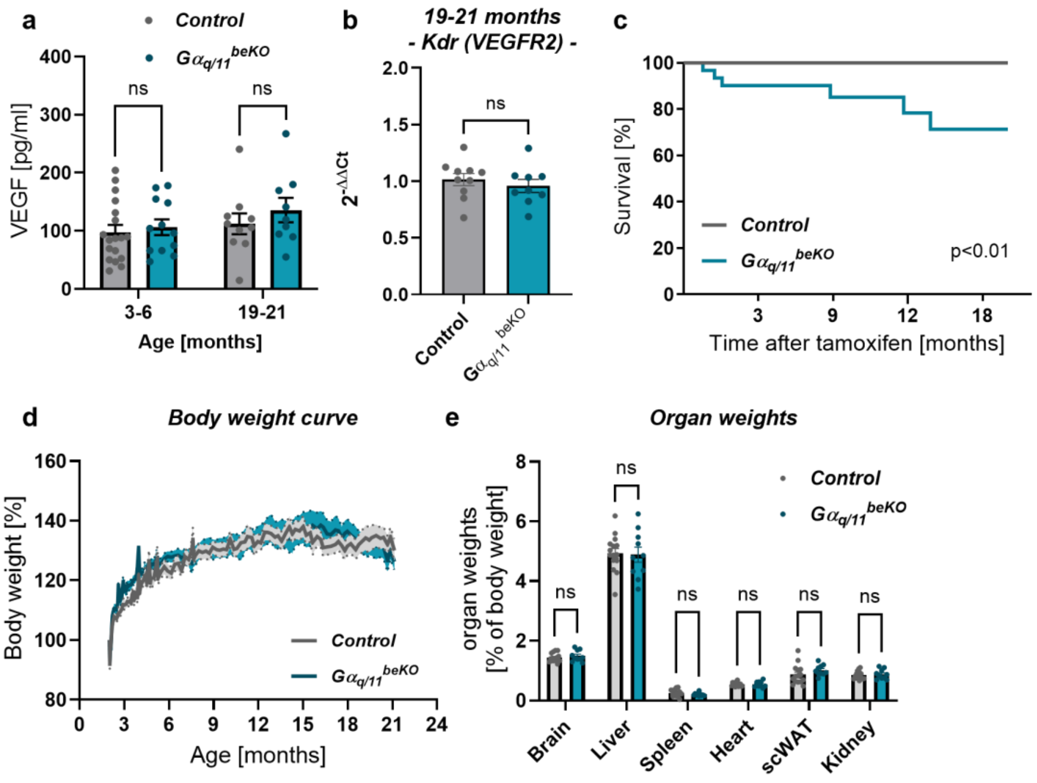

***Supplemental Figure 11: VEGF, survival, and peripheral organs in aged $G\alpha_{q/11}^{beKO}$ mice.*** (**a**) Quantification of VEGF concentration in plasma of 3-6-month-old and 20-22-month-old control and $G\alpha_{q/11}^{beKO}$ mice. N=9-18. (**b**) VEGFR2 mRNA (*Kdr*) expression in brain tissue from 20-22-month-old control and $G\alpha_{q/11}^{beKO}$ mice. (**c**) Survival curve of control and $G\alpha_{q/11}^{beKO}$ mice. Tamoxifen was administered at the age of 7-8 weeks. N=28-32. Log-rank (Mantel-Cox) test. (**d**) Body weight curve of control and $G\alpha_{q/11}^{beKO}$ mice. Tamoxifen was administered at the age of 7-8 weeks. (**e**) Organ weights as percentage of body weight of control and $G\alpha_{q/11}^{beKO}$ mice at the age of 20-22 months. N=10-15. Shown are means±SEM. ns: $p>0.05$. Detailed information on the age and time after tamoxifen, as well as the exact test statistics and values, is provided in Supplemental Table 3.

SUPPLEMENTAL METHODS

*Carotid artery preparation*

To assess the recombination in carotid arteries, the mice were perfused with 4% paraformaldehyde after Ringer perfusion under deep anaesthesia. Subsequently, the region where the common carotid artery diverges into the internal and the external carotid arteries was prepared under a stereomicroscope. Then, a tissue block was removed and snap-frozen. Cryosections (20 µm) were prepared using a cryostat (CM3050, Leica) and costained with DAPI (#A1001, BioChemica, 1 µg/mL). The images were taken using a fluorescence microscope (DMI6000B, Leica).

*Arterial Spin Labelling MRI measurements*

Anaesthesia was induced by 4% and maintained at 0.8-1% isoflurane in 21% $O_2$ and 79% $N_2$. The respiration rate and body temperature were continuously monitored using an MRI-compatible system, and the temperature was maintained at 37±0.2°C throughout the study using a feedback-regulated warming system (Small Animal Instruments Inc.). The mice were scanned using a 7 Tesla small animal MRI scanner (ClinScan, Bruker) with a $^{1}H$ receive-only 2x2 mouse brain surface array coil. The array coil was placed close to the neck and back of the head. Anatomical image data were acquired using a T2-weighted turbo spin-echo sequence with the following sequence parameters: echo time – TE = 65 ms, repetition time – TR = 3000 ms, field of view – FoV = 16×16 $mm^2$, image matrix = 192×192, turbo factor = 9, number of slices = 15, slice thickness = 0.5 mm, readout bandwidth = 100 Hz/pixel, 50% phase oversampling, and total acquisition time = 3:15 min. CBF was measured by arterial spin labelling (ASL) with Q2TIPS (quantitative imaging of perfusion using a single subtraction with interleaved thin-slice TI1 periodic saturation)(3) with TI1/TI2/SS = 900/1400/1375 ms and with 45 control-label pairs. Other ASL scanning parameters were: Echo planar imaging technique with TE =7.3 ms, TR =3000 ms, FoV =16×16 $mm^2$, image matrix = 64×64, number of slices = 3, slice thickness = 0.5 mm, partial Fourier factor =6/8, readout bandwidth = 4340 Hz/pixel, and total acquisition time = 4:38 min.

*Time-of-Flight (TOF) MR angiography*

Time-of-Flight (TOF) MR angiography was performed during the same imaging session as follows: TE = 3.06 ms, TR = 18 ms, FoV = 20×20 $mm^2$, image matrix = 256×256, partial Fourier factor in phase encoding direction = 6/8, number of imaging slabs = 4, number of slices per slab = 28, slab overlap = 18%, slice thickness = 0.13 mm, readout bandwidth = 151 Hz/pixel, total acquisition time = 6:24 min. The voxel size of the data sets was 78x78x51 µm. For analyses of the cerebral arterial vasculature, the TOF data sets were analysed using IMARIS (v9.9.1, Bitplane) as illustrated in Supplemental Fig. 5. An autopath tracing algorithm was used for the analysis of the middle cerebral artery (MCA) and posterior cerebral artery (PCA) of the left and right hemispheres. In brief, we used the surface creation feature of the software, followed by an automatic thresholding adjustment to subtract background signals, and a manual deletion of signals that were not part of the cerebral artery network. A mask of the final surface was created, and the filament tracing tool of the software was used to generate an autopath. To analyse only the vessel structure of the MCAs and PCAs, the starting points were set to the diverging points at the Circle of Willis. The mean diameter, the volumes, and the straightness were measured in all detectable branches of the traced MCAs and PCAs and averaged for each animal.

*Real-time RT-PCR*

RNA was isolated from brain tissue, PBECs, or FACS-isolated brain endothelial cells by Nucleo Spin RNA Kit (Macherey-Nagel) according to the manufacturer´s instructions. RNA (similar amount of each sample) was transcribed with Moloney Murine Leukemia Virus Reverse Transcriptase and random hexamer primers (Promega). The following primers were used for quantitative real-time PCR: *Ppia* forward, 5´-GCATACAGGTCCTGGCATCT-3´, *Ppia* reverse, 5´-CATCCAGCCATTCAGTCTTGG-3´; *Gnaq* forward, 5’-GGTTCCAGAACTCCTCTGTG-3’, *Gnaq* reverse, 5’-CTCTCTGGGGTCCATCATATTCT-3’; *Gna11* forward, 5’-ATCAAGACGCTGTGGAGTGA-3’, *Gna11* reverse, 5’-GTCCACGTCCGTCAAGTAGT-3’; *Cdkn1a* forward, 5’-CCTGGTGATGTCCGACCTG-3’, *Cdkn1a* reverse, 5’-CCATGAGCGCATCGCAATC-3’; *Kdr* forward, 5’-TTTGGCAAATACAACCCTTCAGA-3’, *Kdr* reverse, 5’-GCTCCAGTATCATTTCCAACCA-3’; *Tjp1* forward, 5’-GGAGATGTTTATGCGGACGGT-3’, *Tjp1* reverse, 5’-TCCTCCATTGCTGTGCTCTTAG-3’; *Ocln* forward, 5’-TACTGGTCTCTACGTGGATCAAT-3’, *Ocln* reverse, 5’-TTCTTCGGGTTTTCACAGCAA-3’. Quantitative RT-PCR was performed according to the following protocol: after a 10 min preincubation at 95 °C, amplification was performed for 30 sec at 95 °C, 30 sec at 60 °C, 30 sec at 72 °C, and 1 min at 60 °C (38 - 42 cycles). Amplification was quantified using Platinum SYBR Green qPCR SuperMix (Invitrogen). Quantified results were normalised to *Ppia* and the respective control group using the ΔΔCt method.

*Fluorescence-activated cell sorting of brain endothelial cells*

For the isolation of single endothelial cells of mouse brains of different ages, we performed preparation and sorting as described previously.(4) Briefly, after decapitation, we homogenised a half hemisphere in dissection medium (15 mmol/L HEPES, 3.6 mg/mL glucose in HBSS), and incubated the tissue in 5 mg/mL Collagenase/Dispase (Roche Diagnostics), 0.02 mg/mL DNAse I (Roche Diagnostics), 0.5 mg/mL TLCK (Sigma-Aldrich) at 37 °C for 30 minutes. After filtering the cells through a 70-µm strainer, they were separated from myelin by gradient centrifugation in 37% Percoll (800 x *g*, 30 min, 4 °C). Cells were stained with Live/Dead staining (No. 65-0865-14, Thermo Fisher, 1:2000), and anti-mouse CD31 and anti-CD45 antibodies for 45 minutes on ice after incubating in PBS containing FC block (anti-mouse CD16/CD32, No. 553142, BD Pharmingen, 2.5 µg/mL) for 15 min. Cells were sorted using a FACSAria III cell sorter (BD Biosciences) and lysed in RNA lysis buffer as described above.

*VEGF measurement*

The concentration of VEGF was determined in blood collected from the heart of deeply anaesthetised mice. After centrifuging at 16,400 x *g* and 4 °C for five minutes, EDTA plasma was stored at -80 °C before measurement. The amount of VEGF was determined by using a commercially available ELISA kit (RAB0509, Sigma-Aldrich) according to the manufacturer's protocol.

SUPPLEMENTAL TABLES

***Table 1. Primary antibodies used for Western blotting, immunofluorescence stainings, and fluorescence-activated cell sorting (FACS).***

| **Target** | **Company (catalogue no.)** | **Dilution** | **RRID** | **Application** |
|---|---|---|---|---|
| 4-HNE | Merck Millipore (393207) | 1:200 | AB_566310 | Immunofluorescence |
| Actin | Millipore Sigma (MAB1501) | 1:2500 | AB_2223041 | Western blotting |
| AKT | Cell Signaling Technology (9272) | 1:1000 | AB_329827 | Western blotting |
| Albumin | Bethyl (A90-134) | 1:5000 | AB_67120 | Western blotting |
| Bassoon | Synaptic Systems (141016) | 1:500 | | Immunofluorescence |
| Caveolin 1 | Cell Signaling Technology (3267) | 1:100 | AB_2275453 | Immunofluorescence |
| CD13 | AbD Serotec (MCA2183GA) | 1:200 | AB_323548 | Immunofluorescence |
| CD31 | Bio-Rad (MCA2388) | 1:100 | AB_2161026 | Immunofluorescence |
| CD31 (APC-coupled) | MEC13.3; BioLegend (102509) | 1:100 | AB_312916 | FACS |
| CD45 (RB705-coupled | 30-F11 BD Biosciences (570291) | 1:100 | AB_3086726 | FACS |
| Claudin 5 | Invitrogen (34-1600) | 1:400 | AB_2533157 | Immunofluorescence |
| Collagen IV | Bio-Rad (134001) | 1:200 | AB_2082646 | Immunofluorescence |
| eNOS | BD Transduction (610296) | 1:2000 | AB_397690 | Western blotting |
| ERK1/2 | Cell Signaling Technology (4695) | 1:1000 | AB_390779 | Western blotting |
| GFAP | Millipore (AB5541) | 1:400 | AB_177521 | Immunofluorescence |
| Iba1 | Fujifilm Wako (019-19741) | 1:400 | AB_839504 | Immunofluorescence |
| Lamin B1 | Proteintech (12987-1-AP) | 1:100 | AB_2136290 | Immunofluorescence |
| Laminin α5 | customised by L. Sorokin, Münster | 1:100 | Self-made | Immunofluorescence |
| MBP | Millipore (MAB386) | 1:250 | AB_94975 | Immunofluorescence |
| NeuN | Millipore (MAB377) | 1:500 | AB_2298772 | Immunofluorescence |
| pAKT | Cell Signaling Technology (4060) | 1:2000 | AB_2315049 | Western blotting |
| peNOS | Cell Signaling Technology (9571) | 1:1000 | AB_329837 | Western blotting |
| pERK1/2 | Cell Signaling Technology (9101) | 1:1000 | AB_331646 | Western blotting |
| pTau AT180 (Thr231) | Thermo Fischer Scientific (MN1040) | 1:500 | AB_223649 | Western blotting |
| pTau AT8 (Ser202/Thr205) | Thermo Fischer Scientific (MN1020) | 1:1000 | AB_223647 | Western blotting |
| pVEGFR2 | Cell Signaling Technology (2478) | 1:1000 | AB_331377 | Western blotting |
| pγH2AX | Cell Signaling Technology (9718T) | 1:400 | AB_2118009 | Immunofluorescence |
| Shank2 | Synaptic Systems (162204) | 1:500 | AB_2619861 | Immunofluorescence |
| SMA | Sigma Aldrich (C6198) | | AB_476856 | Immunofluorescence |
| Tau | Cell Signaling Technology (4019S) | 1:1000 | AB_10695394 | Western blotting |
| Tubulin | Cell Signaling Technology (2144) | 1:1000 | AB_2210548 | Western blotting |
| VEGFR$_2$ | Cell Signaling Technology (2479) | 1:1000 | AB_2212507 | Western blotting |

***Supplemental Table 2. Secondary antibodies used for Western blotting and immunofluorescence stainings.***

| **Target (coupled to)** | **Company (catalogue no.)** | **Dilution** | **RRID** | **Application** |
|---|---|---|---|---|
| Chicken IgY (Alexa Fluor 488) | Abcam (ab150169) | 1:200 | AB_2636803 | Immunofluorescence |
| Goat IgG (Alexa Fluor 488) | Thermo Fisher (A-11055) | 1:200 | AB_2534102 | Immunofluorescence |
| Goat IgG (Alexa Fluor 555) | Thermo Fisher (A-21432) | 1:200 | AB_2535853 | Immunofluorescence |
| Goat IgG (HRP) | DakoCytomation (P0160) | 1:2500 | AB_656964 | Western blotting |
| Guinea pig (Alexa Fluor 633) | Thermo Fisher (A-21105) | 1:400 | AB_2535757 | Immunofluorescence |
| Mouse (Alexa Fluor 555) | Thermo Fisher (A-21422) | 1:400 | AB_141822 | Immunofluorescence |
| Mouse IgG (HRP) | DakoCytomation (P0447) | 1:2500 | AB_2617137 | Western blotting |
| Rabbit IgG (Alexa Fluor 488) | Thermo Fisher (A-21206) | 1:200 | AB_2535792 | Immunofluorescence |
| Rabbit IgG (Alexa Fluor 555) | Thermo Fisher (A-31572) | 1:200 | AB_162543 | Immunofluorescence |
| Rabbit IgG (Alexa Fluor 750) | Thermo Fisher (A-21039) | 1:400 | AB_2535710 | Immunofluorescence |
| Rabbit IgG (HRP) | DakoCytomation (P0448) | 1:2500 | AB_2617138 | Western blotting |
| Rat IgG (Alexa Fluor 647) | Abcam (ab150155) | 1:400 | AB_2813835 | Immunofluorescence |
| Rat IgG (Alexa Fluor 488) | Thermo Fisher (A-21208) | 1:200 | AB_2535794 | Immunofluorescence |
| Rat IgG (Alexa Fluor 555) | Abcam (ab150154) | 1:200 | AB_2813834 | Immunofluorescence |

***Supplemental Table 3. Results of statistical analyses and age information***

| Figure | Age of the mice | Time after tamoxifen | Sex | Statistical test | Values |
|---|---|---|---|---|---|
| 1b, c | 3-6 months | 2-5 weeks | 7 female, 8 male | 1c: Two-tailed Mann-Whitney test | p=0.0140 |
| 1e | 3-6 months | 2-5 weeks | 9 female, 11 male | Repeated measures Two-Way ANOVA followed by Sidak´s post-hoc tests (p-values adjusted for multiple comparisons) | **Time**: F(80,1440)=61.93, p<0.0001; **Genotype**: F(1,18)=4.064, p=0.059; **Interaction**: F(80,1440)=2.626, p<0.0001 |
| 1f | 3-6 months | 2-5 weeks | 9 female, 11 male | Two-tailed unpaired t test | p=0.0388 |
| 1h | 3-6 months | 2-5 weeks | 7 female, 8 male | Two-tailed unpaired t test | p=0.8037 |
| 1i, j | 2-11 months | 2-4 weeks | 5 female, 6 male | 1j: Two-tailed Mann-Whitney test | p=0.0268 |
| 2c | 3-5 months | 2-3 weeks | 14 female, 8 male | Mixed Effects analysis (REML) | **Branch**: F(4,76)=9.809, p<0.0001; **Genotype**: F(1,20)=3.909, p=0.062; **Interaction**: F(4,76)=1.577, p=0.1890 |
| 2d | 3-5 months | 2-3 weeks | 14 female, 8 male | Repeated measures Two-Way ANOVA | **Branch**: F(4,80)=6.379, p=0.0002; **Genotype**: F(1,20)=1.293, p=0.2689; **Interaction**: F(4,80)=2.637, p=0.0399 |
| 2e | 3-5 months | 2-3 weeks | 14 female, 8 male | Mixed Effects analysis (REML) | **Branch**: F(4,76)=7.706, p<0.0001; **Genotype**: F(1,20)=0.1359, p=0.7163; **Interaction**: F(4,76)=1.452, p=0.2251 |
| 2f | 3-5 months | 2-3 weeks | 14 female, 8 male | Repeated measures Two-Way ANOVA | **Branch**: F(4,80)=7.350, p<0.0001; **Genotype**: F(1,20)=5.302, p=0.0322; **Interaction**: F(4,80)=0.5599, p=0.6924 |
| 2g | 3-5 months | 2-3 weeks | 13 female, 8 male | Repeated measures Two-Way ANOVA | **Branch**: F(4,76)=2.424, p=0.0553; **Genotype**: F(1,19)=0.2444, p=0.6267; **Interaction**: F(4,76)=2.349, p=0.0617 |
| 2h | 3-5 months | 2-3 weeks | 14 female, 8 male | Repeated measures Two-Way ANOVA | **Branch**: F(4,80)=1.575, p=0.1889; **Genotype**: F(1,20)=5.046, p=0.0361; **Interaction**: F(4,80)=1.073, p=0.3756 |
| 2j | 3-5 months | 2-5 weeks | 4 female, 5 male | Repeated measures Two-Way ANOVA | **Depth:** F(4,28)=30.03, p<0.0001; **Genotype:** F(1,7)=0.0087, p=0.9281; **Interaction:** F(4,28)=0.2521, p=0.9095 |
| 2k | 3-5 months | 2-5 weeks | 4 female, 5 male | Repeated measures Two-Way ANOVA | **Depth:** F(4,28)=23.69, p<0.0001; **Genotype:** F(1,7)=6.129, p=0.0425; **Interaction:** F(4,28)=0.0423, p=0.9964 |
| 2l | 3-5 months | 2-5 weeks | 4 female, 5 male | Two-tailed unpaired t test | p=0.9755 |
| 2m | 3-5 months | 2-5 weeks | 4 female, 4 male | Repeated measures Two-Way ANOVA | **Vessel type**: F(2,18)=60.53, p<0.0001; |

| | | | | | |
|---|---|---|---|---|---|
| | | | | followed by Sidak´s post-hoc tests (p-values adjusted for multiple comparisons) | **Genotype**: $F(1,18)=0.9112$, $p=0.3524$; **Interaction**: $F(2,18)=0.0058$, $p=0.9942$ |
| 2o | 3-6 months | 2-5 weeks | 14 female, 12 male | Repeated measures Two-Way ANOVA followed by Sidak´s post-hoc tests (p-values adjusted for multiple comparisons) | **Brain region**: $F(4,96)=54.03$, $p<0.0001$; **Genotype**: $F(1,24)=0.0383$, $p=0.8466$; **Interaction**: $F(4,96)=1.154$, $p=0.3361$ |
| 3b | 2-5 months | not applicable | 9 female, 10 male | Two-tailed unpaired t test | $p=0.0022$ |
| 3c | 2-5 months | not applicable | 9 female, 10 male | Two-tailed unpaired t test | $p=0.0686$ |
| 3d | 4-8 months | not applicable | 6 female, 4 male | Two-tailed unpaired t test | $p=0.0199$ |
| 3f, g | 7-8 months | not applicable | 4 female | 3g: Two-tailed Mann-Whitney test | $p=0.0286$ |
| 3h | 5-7 months | 3 months | 5 female, 10 male | Two-tailed unpaired t test | $p=0.0019$ |
| 3i | 5-7 months | 3 months | 5 female, 10 male | Two-tailed Mann-Whitney test | $p=0.0176$ |
| 3j | 5-7 months | 3 months | 5 female, 10 male | Two-tailed unpaired t test | $p=0.0083$ |
| 3k | 5-7 months | 3 months | 5 female, 10 male | Two-tailed unpaired t test | $p=0.0305$ |
| 4b | 2 months | not applicable | 18 female, 19 male | One sample t test to 0.5 chance level | Control: $p=0.0020$ $G\alpha_{q/11}^{beKO}$: $p=0.0004$ |
| 4c | 6 months | 4 months | 16 female, 17 male | One sample t test to 0.5 chance level | Control: $p=0.0013$ $G\alpha_{q/11}^{beKO}$: $p=0.0355$ |
| 4d | 12 months | 10 months | 16 female, 17 male | One sample t test to 0.5 chance level | Control: $p<0.0001$ $G\alpha_{q/11}^{beKO}$: $p=0.0031$ |
| 4e | 18 months | 16 months | 12 female, 15 male | One sample t test to 0.5 chance level | Control: $p=0.0054$ $G\alpha_{q/11}^{beKO}$: $p=0.6857$ |
| 4g-i | 20-22 months | 18-20 months | 12 female, 15 male | 4i: Scheirer-Ray-Hare test followed by Mann-Whitney post hoc tests, Bonferroni-Sidak correction | **Genotype**: $\chi 2(1)=10.85$, $p=0.0044$; |
| 4j | 20-22 months | 18-20 months | 12 female, 15 male | Repeated measures Two-Way ANOVA | **Time**: $F(4,100)=37.59$, $p<0.0001$; **Genotype**: $F(1,25)=4.542$, $p=0.0431$; **Interaction**: $F(4,100)=1.726$, $p=0.1501$ |
| 4k | 20-22 months | 18-20 months | 12 female, 15 male | Mixed Effects analysis (REML) followed by Sidak´s post-hoc tests (p-values adjusted for multiple comparisons) | **Time**: $F(1,25)=0.3478$, $p=0.5607$; **Genotype**: $F(1,26)=5.502$, $p=0.0269$; **Interaction**: $F(1,25)=3.018$, $p=0.0946$ |
| 4l | 20-22 months | 18-20 months | 12 female, 14 male | Mixed Effects analysis (REML) followed by Sidak´s post-hoc tests (p-values adjusted for multiple comparisons) | **Time**: $F(1,24)=0.3.903$, $p=0.0598$; **Genotype**: $F(1,25)=3.448$, $p=0.0752$; **Interaction**: $F(1,24)=6.200$, $p=0.0201$ |
| 5b | 20-22 months | 18-20 months | 12 female, 14 male | Two-Way ANOVA followed by Sidak´s post-hoc tests (p-values adjusted for multiple comparisons) | **Brain region**: $F(1,48)=33.86$, $p<0.0001$; **Genotype**: $F(1,48)=0.1140$, $p=0.7371$; **Interaction**: $F(1,48)=0.2622$, $p=0.6109$ |

| | | | | | |
|---|---|---|---|---|---|
| 5d | 20-22 months | 18-20 months | 12 female, 14 male | Two-Way ANOVA followed by Sidak´s post-hoc tests (p-values adjusted for multiple comparisons) | **Brain region**: F(1,48)=5.012, p=0.0298; **Genotype**: F(1,48)=4.344, p=0.0425; **Interaction**: F(1,48)=0.5390, p=0.4664 |
| 6b | 20-22 months | 18-20 months | 10 female, 10 male | Scheirer-Ray-Hare test followed by Mann-Whitney post hoc tests, Bonferroni-Sidak correction | **Brain region**: $\chi 2(2)$ =5.62, p=0.060; **Genotype**: $\chi 2(1)$ =9.00, p=0.003; **Interaction**: $\chi 2(2)$=0.06, p=0.971 |
| 6c | 20-22 months | 18-20 months | 10 female, 10 male | Two-tailed unpaired t test | p=0.0108 |
| 6d (left) | 4-6 months | 4 weeks | 10 female, 12 male | Two-tailed unpaired t test | p=0.9497 |
| 6d (right) | 13-14 months | 11-12 months | 5 female, 5 male | Two-tailed unpaired t test | p=0.8705 |
| 6e | 20-22 months | 18-20 months | 9 female, 10 male | Two-tailed Mann-Whitney test | p=0.0028 |
| 6g | 20-22 months | 18-20 months | 10 female, 10 male | Repeated measures Two-Way ANOVA followed by Sidak´s post-hoc tests (p-values adjusted for multiple comparisons) | **Brain region**: F(1,18)=16.81, p=0.0007; **Genotype**: F(1,18)=7.841, p=0.0118; **Interaction**: F(1,18)=0.0007, p=0.9789 |
| 6i | 20-22 months | 18-20 months | 10 female, 10 male | Repeated measures Two-Way ANOVA followed by Sidak´s post-hoc tests (p-values adjusted for multiple comparisons) | **Brain region**: F(1,18)=0.088, p=0.7700; **Genotype**: F(1,18)=5.041, p=0.0375; **Interaction**: F(1,18)=3.707, p=0.0701 |
| 6j | 20-22 months | 18-20 months | 10 female, 10 male | Scheirer-Ray-Hare test followed by Mann-Whitney post hoc tests, Bonferroni-Sidak correction | **Brain region**: $\chi 2(1)$ =1.08, p=0.299; **Genotype**: $\chi 2(1)$ =4.16, p=0.041; **Interaction**: $\chi 2(1)$=0.05, p=0.823 |
| 7a | 20-22 months | 18-20 months | 10 female, 10 male | Two-tailed unpaired t test | p=0.0248 |
| 7c, d | 7-8 months | not applicable | 4 female | 7d: Two-tailed unpaired t test | p=0.0127 |
| 7f | 7 months | 3 weeks | 3 male | Repeated measures Two-Way ANOVA followed by Sidak´s post-hoc tests (p-values adjusted for multiple comparisons) | **Treatment**: F(1,4)=130.5, p=0.0003; **Genotype**: F(1,4)=0.0254, p=0.8811; **Interaction**: F(1,4)=0.0260, p=0.8798 |
| 7g | 7 months | 3 weeks | 3 male | Repeated measures Two-Way ANOVA followed by Sidak´s post-hoc tests (p-values adjusted for multiple comparisons) | **Treatment**: F(1,4)=16.74, p=0.0150; **Genotype**: F(1,4)=0.0294, p=0.8722; **Interaction**: F(1,4)=0.0237, p=0.8854 |
| 7h | 7 months | 3 weeks | 3 male | Repeated measures Two-Way ANOVA followed by Sidak´s post-hoc tests (p-values adjusted for multiple comparisons) | **Treatment**: F(1,4)=8.515, p=0.0433; **Genotype**: F(1,4)=9.371, p=0.0376; **Interaction**: F(1,4)=5.957, p=0.0712 |
| 7i | 7 months | 3 weeks | 3 male (6 samples) | Two-tailed unpaired t test | p=0.1596 |

| | | | | | |
|---|---|---|---|---|---|
| S1a left | 3-5 months | 2 weeks | 4 female, 8 male | Two-tailed Mann-Whitney test | p=0.0022 |
| S1a right | 3-5 months | 2 weeks | 4 female, 8 male | Two-tailed Mann-Whitney test | p=0.0022 |
| S1b | 2-5 months | 2-3 weeks | 7 female, 6 male | Two-tailed Mann-Whitney test | p=0.0012 |
| S2a-b | 3-6 months | 2-5 weeks | 10 female, 14 male | S2b: Two-tailed Mann-Whitney test | p=0.5195 |
| S2c-d | 3-4 months | 2-3 weeks | 7 female, 7 male | S2d: Two-tailed Mann-Whitney test | p=0.9372 |
| S3a left | 5-6 months | 10-12 weeks after AAV injection | 5 female, 8 male | Two-tailed unpaired t test | p=0.0164 |
| S3a right | 5-6 months | 10-12 weeks after AAV injection | 5 female, 8 male | Two-tailed Mann-Whitney test | p=0.0022 |
| S3b, c | 5-6 months | 10-12 weeks after AAV injection | 10 female, 16 male | S3c: Two-tailed unpaired t test | p=0.0393 |
| S4 | 4 months | 4 weeks | 1 male | | |
| S6a | 18 months | 16 months | 12 female, 15 male | Two-tailed Mann-Whitney test | p=0.0743 |
| S6a | 18 months | 16 months | 12 female, 15 male | Two-tailed Mann-Whitney test | p=0.0743 |
| S6b | 20-22 months | 18-20 months | 12 female, 15 male | Two-tailed Mann-Whitney test | p=0.0743 |
| S6c | 20-22 months | 18-20 months | 12 female, 15 male | Mixed Effects analysis (REML) followed by Sidak´s post-hoc tests (p-values adjusted for multiple comparisons) | **Time**: F(1,24)=0.5037, p=0.4847; **Genotype**: F(1,25)=8.382, p=0.0078; **Interaction**: F(1,24)=2.689, p=0.1135 |
| S6d | 20-22 months | 18-20 months | 12 female, 15 male | Mixed Effects analysis (REML) followed by Sidak´s post-hoc tests (p-values adjusted for multiple comparisons) | **Time**: F(1,24)=0.1289, p=0.7228; **Genotype**: F(1,25)=2.610, p=0.1187; **Interaction**: F(1,24)=1.142, p=0.2959 |
| S7c | 20-22 months | 18-20 months | 11 female, 14 male | Two-tailed unpaired t test | p=0.5360 |
| S7d | 20-22 months | 18-20 months | 11 female, 14 male | Two-tailed unpaired t test | p=0.7013 |
| S7e | 20-22 months | 18-20 months | 11 female, 14 male | Two-tailed unpaired t test | p=0.8938 |
| S7f | 20-22 months | 18-20 months | 11 female, 14 male | Two-tailed unpaired t test | p=0.7418 |
| S7g | 20-22 months | 18-20 months | 11 female, 14 male | Two-tailed unpaired Mann Whitney test | p=0.8460 |
| S7h | 20-22 months | 18-20 months | 11 female, 14 male | Two-tailed unpaired t test | p=0.8232 |
| S7i | 20-22 months | 18-20 months | 11 female, 14 male | Two-tailed unpaired t test | p=0.7988 |
| S7j | 20-22 months | 18-20 months | 11 female, 14 male | Two-tailed unpaired t test | p=0.5816 |
| S7k | 20-22 months | 18-20 months | 11 female, 14 male | Two-tailed unpaired t test | p=0.4258 |
| S8b | 20-22 months | 18-20 months | 11 female, 15 male | Two-tailed unpaired t test | p=0.4054 |

| | | | | | |
|---|---|---|---|---|---|
| S8c | 20-22 months | 18-20 months | 11 female, 15 male | Two-tailed unpaired Mann Whitney test | p=0.6832 |
| S8d | 20-22 months | 18-20 months | 11 female, 15 male | Two-tailed unpaired Mann Whitney test | p=0.2372 |
| S8f | 20-22 months | 18-20 months | 11 female, 15 male | Two-tailed unpaired t test | p=0.3289 |
| S8g | 20-22 months | 18-20 months | 11 female, 15 male | Two-tailed unpaired t test | p=0.5557 |
| S8h | 20-22 months | 18-20 months | 11 female, 15 male | Two-tailed unpaired t test | p=0.2339 |
| S8i | 20-22 months | 18-20 months | 11 female, 15 male | Two-tailed unpaired t test | p=0.1105 |
| S9b | 6-20 months | not applicable | 10 female, 2 male | One-Way ANOVA | F(2,9)=0.01693 p=0.9832 |
| S9c | 6-20 months | not applicable | 9 female, 2 male | One-Way ANOVA | F(2,8)=0.3632 p=0.7064 |
| S10a | 20-22 months | 18-20 months | 10 female, 10 male | Two-tailed unpaired t test | p=0.6927 |
| S10b | 20-22 months | 18-20 months | 9 female, 9 male | Two-tailed unpaired t test | p=0.4499 |
| S10c | 20-22 months | 18-20 months | 11 female, 15 male | Two-tailed unpaired t test | p=0.2228 |
| S10d | 20-22 months | 18-20 months | 11 female, 14 male | Two-tailed unpaired Mann Whitney test | p=0.1289 |
| S10e | 20-22 months | 18-20 months | 10 female, 10 male | Repeated measures Two-Way ANOVA followed by Sidak´s post-hoc tests (p-values adjusted for multiple comparisons) | **Brain region**: F(1,18)=119.1, p<0.0001; **Genotype**: F(1,18)=3.489, p=0.0781; **Interaction**: F(1,18)=2.220, p=0.1535 |
| S10f | 13-14 months | 11-12 months | 7 female, 6 male | Two-tailed unpaired t test | p=0.9252 |
| S10g | 20-22 months | 18-20 months | 10 female, 10 male | Repeated measures Two-Way ANOVA followed by Sidak´s post-hoc tests (p-values adjusted for multiple comparisons) | **Brain region**: F(1,18)=216.6, p<0.0001; **Genotype**: F(1,18)=1.708, p=0.2077; **Interaction**: F(1,18)=0.0388, p=0.8461 |
| S10h | 4-6 months | 4 weeks | 6 female, 7 male | Mixed Effects analysis (REML) followed by Sidak´s post-hoc tests (p-values adjusted for multiple comparisons) | **Brain region**: F(1,10)=5.095, p=0.0476; **Genotype**: F(1,11)=0.649, p=0.4374; **Interaction**: F(1,10)=0.2387, p=0.6357 |
| S10i | 4-6 months | 4 weeks | 6 female, 7 male | Repeated measures Two-Way ANOVA followed by Sidak´s post-hoc tests (p-values adjusted for multiple comparisons) | **Brain region**: F(1,11)=0.0017, p=0.9672; **Genotype**: F(1,11)=0.2206, p=0.6477; **Interaction**: F(1,11)=4.818, p=0.0505 |
| S10j | 6-8 months | 4 months | 8 female, 4 male | Repeated measures Two-Way ANOVA followed by Sidak´s post-hoc tests (p-values adjusted for multiple comparisons) | **Brain region**: F(1,10)=0.4013, p=0.5406; **Genotype**: F(1,10)=4.199, p=0.0676; **Interaction**: F(1,10)=0.4013, p=0.5406 |
| S10k | 13-14 months | 11-12 months | 7 female, 6 male | Mixed Effects analysis (REML) followed by Sidak´s post-hoc tests (p-values adjusted for multiple comparisons) | **Brain region**: F(1,10)=1.443, p=0.2574; **Genotype**: F(1,11)=0.7176, p=0.4150; |

| | | | | | |
|---|---|---|---|---|---|
| | | | | | **Interaction**: F(1,10)=1.443, p=0.2574 |
| S10l | 20-22 months | 18-20 months | 8 female, 10 male | Two-tailed unpaired Mann Whitney test | p=0.0789 |
| S10m | 20-22 months | 18-20 months | 9 female, 10 male | Two-tailed unpaired Mann Whitney test | p=0.6185 |
| S10n | 20-22 months | 18-20 months | 8 female, 10 male | Two-tailed unpaired t test | p=0.9230 |
| S11a | 3-6 months and 20-22 months | 3-5 weeks and 18-20 months | 22 female, 27 male | Two-Way ANOVA followed by Sidak´s post-hoc tests (p-values adjusted for multiple comparisons) | **Age**: F(1,45)=1.887, p=0.1763; **Genotype**: F(1,45)=1.037, p=0.3139; **Interaction**: F(1,45)=0.2265, p=0.6365 |
| S11b | 20-22 months | 18-20 months | 9 female, 10 male | Two-tailed unpaired t test | p=0.4864 |
| S11c | 2-22 months | 0-20 months | 27 female, 33 male | Survival analysis with Kaplan-Meier followed by Log-rank (Mantel-Cox) test to compare curves | $\chi2(1)$ =7.289 p=0.0069 |
| S11d | 2-22 months | 0-20 months | 18 female, 19 male | no comparisons performed | |
| S11e | 20-22 months | 18-20 months | 11 female, 14 male | Each organ individual two-tailed unpaired t tests; Except spleen: Mann Whitney test | Brain: p=0.4273<br>Liver: p=0.8595<br>Spleen: p=0.3177<br>Heart: p=0.9005<br>scWAT: p=0.1960<br>Kidney: p=0.7351 |

SUPPLEMENTAL REFERENCES

1. Ximerakis M, Lipnick SL, Innes BT, Simmons SK, Adiconis X, Dionne D, et al. Single-cell transcriptomic profiling of the aging mouse brain. *Nature neuroscience.* 2019;22(10):1696-708.
2. Tarhan L, Bistline J, Chang J, Galloway B, Hanna E, and Weitz E. Single Cell Portal: an interactive home for single-cell genomics data. *bioRxiv.* 2023.
3. Luh WM, Wong EC, Bandettini PA, and Hyde JS. QUIPSS II with thin-slice TI1 periodic saturation: a method for improving accuracy of quantitative perfusion imaging using pulsed arterial spin labeling. *Magnetic resonance in medicine.* 1999;41(6):1246-54.
4. Wenzel J, Spyropoulos D, Assmann JC, Khan MA, Stolting I, Lembrich B, et al. Endogenous THBD (Thrombomodulin) Mediates Angiogenesis in the Ischemic Brain-Brief Report. *Arteriosclerosis, thrombosis, and vascular biology.* 2020;40(12):2837-44.
